# A gene desert required for regulatory control of pleiotropic *Shox2* expression and embryonic survival

**DOI:** 10.1101/2020.11.22.393173

**Authors:** Samuel Abassah-Oppong, Brandon J. Mannion, Matteo Zoia, Raquel Rouco, Virginie Tissieres, Cailyn H. Spurrell, Virginia Roland, Fabrice Darbellay, Anja Ljubojevic, Julie Gamart, Tabitha A. Festa-Daroux, Carly S. Sullivan, Eddie Rodríguez-Carballo, Yoko Fukuda-Yuzawa, Riana Hunter, Catherine S. Novak, Ingrid Plajzer-Frick, Stella Tran, Jennifer A. Akiyama, Diane E. Dickel, Javier Lopez-Rios, Iros Barozzi, Guillaume Andrey, Axel Visel, Len A. Pennacchio, John Cobb, Marco Osterwalder

## Abstract

Gene deserts are defined as genomic regions devoid of protein coding genes and spanning more than 500 kilobases, collectively encompassing about 25% of the human genome. Approximately 30% of all gene deserts are enriched for conserved elements with *cis*-regulatory signatures. These are located predominantly near developmental transcription factors (TFs) but despite predicted critical functions, the transcriptional contributions and biological necessity of most gene deserts remain elusive. Here, we explore the *cis*-regulatory impact of a gene desert flanking the *Shox2* gene, a TF indispensable for proximal limb, craniofacial and cardiac pacemaker development. Using a functional genomics approach in mouse embryos we identify the gene desert as a hub for numerous *Shox2*-overlapping enhancers arranged in a globular chromatin domain with tissue-specific features. In accordance, using endogenous CRISPR deletion, we demonstrate that the gene desert interval is essential for *Shox2* transcriptional control in developing limbs, craniofacial compartments, and the heart. Phenotypically, gene desert ablation leads to pacemaker-related embryonic lethality due to *Shox2* depletion in the cardiac sinus venosus. We show that this role is partially mediated through a distal gene desert enhancer, providing evidence for intra-gene desert regulatory robustness. Finally, we uncover a multi-layered functional role of the gene desert by revealing an additional requirement for stylopod morphogenesis, mediated through an array of proximal limb enhancers (PLEs). In summary, our study establishes the *Shox2* gene desert as a fundamental genomic unit that controls pleiotropic gene expression through modular arrangement and coordinated dynamics of tissue-specific enhancers.

## INTRODUCTION

Functional assessment of gene deserts, gene-free chromosomal segments larger than 500 kilobases (kb), has posed considerable challenges since these large noncoding regions were shown to be a prominent feature of the human genome more than 20 years ago^1^. Stable gene deserts (n=172 in the human genome, ∼30% of all gene deserts) share more than 2% genomic sequence conservation between human and chicken, are enriched for putative enhancer elements and frequently located near developmental genes, suggesting a critical role in embryonic development and organogenesis^2–4^. However, genomic deletion of an initially selected pair of gene deserts displayed mild effects on the expression of nearby genes and absence of overt phenotypic alterations^5^. In contrast, gene deserts centromeric and telomeric to the *HoxD* cluster were shown to harbor “regulatory archipelagos” i.e., multiple tissue-specific enhancers that collectively orchestrate spatiotemporal and colinear *HoxD gene* expression in developing limbs and other embryonic compartments^6,7^. These antagonistic gene deserts represent individual topologically associating domains (TADs) separated by the *HoxD* cluster which acts as a dynamic and resilient CTCF-enriched boundary region^8,9^. Despite such critical roles, the functional requirement of only few gene deserts have been studied in detail, including the investigation of chromatin topology and functional enhancer landscapes in the TADs of other developmental key regulators such as *Sox9, Shh* or *Fgf8*^10–12^.

Self-associating TADs identified by 3D chromatin conformation capture are described as primary higher-order chromatin structures that constrain *cis*-regulatory interactions to target genes and facilitate long-range enhancer-promoter (E-P) contacts^13,14^. TADs are thought to emerge through Cohesin-mediated chromatin loop extrusion and are delimited by association of CTCF to convergent binding sites^15,16^. Re-distribution of E-P interactions can lead to pathogenic effects due to perturbation of CTCF-bound TAD boundaries or re-configuration of TADs^10,17^. Therefore, functional characterization of the 3D chromatin topology and transcriptional enhancer landscapes across gene deserts is a prerequisite for understanding the developmental mechanisms underlying mammalian embryogenesis and human syndromes^18^. Recent functional studies in mice have uncovered that mRNA expression levels of developmental regulator genes frequently depend on additive contributions of enhancers within TADs^19–22^. Hereby, the contribution of each implicated enhancer to total gene dosage can vary, illustrating the complexity of transcriptional regulation through E-P interactions^23^. In addition, nucleotide mutations affecting TF binding sites in enhancers can disturb spatiotemporal gene expression patterns, with the potential to trigger phenotypic abnormalities such as congenital malformations due to altered properties of developmental cell populations^24–26^.

In the current study, we focused on the functional characterization of a stable gene desert downstream (centromeric) of the mouse short stature homeobox 2 (*Shox2*) transcription factor (TF). Tightly controlled *Shox2* expression is essential for accurate development of the stylopod (humerus and femur), craniofacial compartments (maxillary-mandibular joint, secondary palate), the facial motor nucleus and its associated facial nerves, and a subset of neurons of the dorsal root ganglia^27–33^. In addition, *Shox2* in the cardiac sinus venosus (SV) is required for differentiation of progenitors of the sinoatrial node (SAN), the dominant pacemaker population during embryogenesis and adulthood^34–36^. *Shox2* inactivation disrupts Nkx2-5 antagonism in SAN pacemaker progenitors and results in hypoplasia of the SAN and venous valves, leading to bradycardia and embryonic lethality^34,35,37^. In accordance with this role, *SHOX2*-associated coding and non-coding variants in humans were implicated with SAN dysfunction and atrial fibrillation^38–40^. Tbx5 and Isl1 cardiac TF regulators were shown to act upstream of *Shox2* in SAN development^41–44^ and *Isl1* is sufficient to rescue *Shox2*-mediated bradycardia in zebrafish hearts^45^.

The human SHOX gene located on the pseudo-autosomal region (PAR1) of the X and Y chromosomes represents a paralog of SHOX2 (on chromosome 3), hence dividing Shox gene function. *SHOX* is associated with defects and syndromes affecting skeletal, limb and craniofacial morphogenesis^28,46,47^. Rodents have lost their *SHOX* gene during evolution along with other pseudo-autosomal genes and mouse *Shox2* features an identical DNA-interacting homeodomain replaceable by human *SHOX* in a mouse knock-in model^27,48^. Remarkably, while *Shox2*/*SHOX2* genes show highly conserved locus architecture, the *SHOX* gene also features a downstream gene desert of similar extension, containing neural (hindbrain) enhancers with overlapping activities^47^. Our previous studies revealed that *Shox2* transcription in the developing mouse stylopod is partially controlled by a pair of human-conserved limb enhancers termed hs741 and hs1262/LHB-A, the latter residing in the gene desert^19,47,49^. However, the rather moderate loss of *Shox2* limb expression in absence of these enhancers indicated increased complexity and potential redundancies in the underlying enhancer landscape^19^.

Here we identified the *Shox2* gene desert as a critical *cis*-regulatory domain encoding an array of distal enhancers with specific subregional activities, predominantly in limb, craniofacial, neuronal, and cardiac cell populations. We found that interaction of these enhancers with the *Shox2* promoter is likely facilitated by a chromatin loop anchored downstream of the *Shox2* gene body and exhibiting tissue-specific features. Genome editing further demonstrated essential pleiotropic functions of the gene desert, including a requirement for craniofacial patterning, limb morphogenesis, and embryonic viability through enhancer-mediated control of SAN progenitor specification. Our results identify the *Shox2* gene desert as a dynamic enhancer hub ensuring pleiotropic and resilient *Shox2* expression as an essential component of the gene regulatory networks (GRNs) orchestrating mammalian development.

## RESULTS

### Gene desert enhancers recapitulate patterns of pleiotropic *Shox2* expression

The gene encoding the *Shox2* transcriptional regulator is located in a 1 megabase (Mb) TAD (chr3:66337001-67337000) and flanked by a stable gene desert spanning 675kb of downstream (centromeric) genomic sequence (**Fig. 1A**). The *Shox2* TAD only contains one other protein coding gene, *Rsrc1*, located adjacent to *Shox2* on the upstream (telomeric) side and known for roles in pre-mRNA splicing and neuronal transcription^50,51^ (**Fig. 1A**). Genes located beyond the TAD boundaries show either near-ubiquitous (*Mlf1*) or *Shox2*-divergent (*Veph1*, *Ptx3*) expression signatures across tissues and timepoints (**Fig. S1A**). While *Shox2* transcription is dynamically regulated in multiple tissues including proximal limbs, craniofacial subregions, cranial nerve, brain, and the cardiac sinus venosus (SV), only a limited number of *Shox2*-associated enhancer sequences have been previously validated in mouse embryos^19,47,49,52^ (**Fig. 1A, B, S1A**). These studies identified a handful of enhancer elements in the *Shox2* TAD driving reporter activity almost exclusively in the mouse embryonic brain (hs1251, hs1262) and limbs (hs741, hs1262, hs638) (Vista Enhancer Browser) (**Fig. 1A**). To predict *Shox2* enhancers more systematically, and to estimate the number of developmental enhancers in the gene desert, we established a map of stringent enhancer activities based on chromatin state profiles^53^ (ChromHMM) and H3K27 acetylation (H3K27ac) ChIP-seq peak calls across 66 embryonic and perinatal tissue-stage combinations from ENCODE^54^ (https://www.encode.project.org) (see ***Methods***). After excluding promoter regions, this analysis identified 20 elements within the *Shox2*-TAD and its border regions, each with robust enhancer marks in at least one of the tissues and timepoints (E11.5-15.5) examined (**Fig. S1A and Table S1**). Remarkably, 17 of the 20 elements mapping to the *Shox2* TAD or border regions were located within the downstream gene desert, with the majority of H3K27ac signatures overlapping *Shox2* expression profiles across multiple tissues and timepoints, indicating a role in regulation of pleiotropic *Shox2* expression (**Figs. 1B, S1A**). The previously validated hs741 and hs1262 limb enhancers were not among these predictions as elevated H3K27ac marks are present at these enhancers already at earlier stages^19,55^.

**Figure 1.**
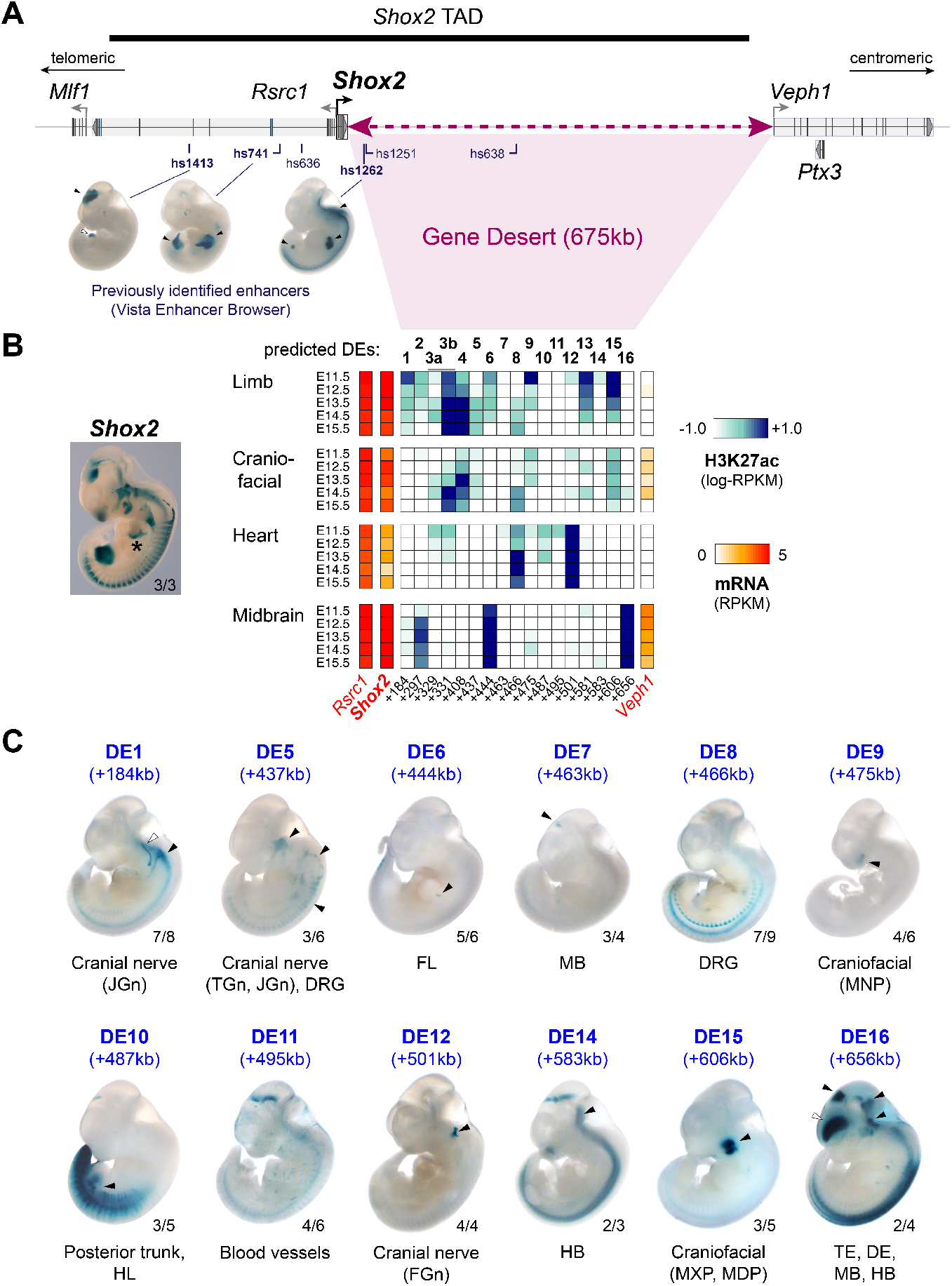
The *Shox2* gene desert constitutes a hub for tissue-specific enhancers. (**A**) Genomic interval containing the *Shox2* TAD^61^ and previously identified *Shox2*-associated enhancer regions (see Vista Enhancer browser). Vista IDs in bold mark enhancers with *Shox2*-overlapping and highly reproducible activities. Hs, homo sapiens. (**B**) Heatmap showing H3K27 acetylation (ac) -predicted and ChromHMM-filtered putative enhancers and their temporal signatures in tissues with dominant *Shox2* functions (see full matrix in **Fig. S1**). Blue and red shades represent H3K27ac enrichment and mRNA expression levels, respectively. Distance to *Shox2* TSS (+) is indicated in kb. Left: *Shox2* expression pattern (*Shox2*-LacZ/+) at E11.5^30^. *, indicates removal of forelimb for better visibility of the heart. (**C**) Transgenic LacZ reporter validation of predicted gene desert enhancers (DEs) in mouse embryos at E11.5. Arrowheads points to reproducible enhancer activity with (black) or without (white) *Shox2* overlap. JGn, TGn, FGn: jugular, trigeminal, and facial ganglion, respectively. DRG, dorsal root ganglia. FL, Forelimb. HL, Hindlimb. TE, Telencephalon. DiE, Diencephalon. MB, Midbrain. HB, Hindbrain. MNP, medial nasal process. MXP, maxillary process. MDP, mandibular process. Reproducibility numbers are indicated on the bottom right of each representative embryo shown (reproducible tissue-specific staining vs. number of transgenic embryos with any LacZ signal).

To determine the *in vivo* activity patterns for each of the predicted gene desert enhancer (DE) elements, we performed *LacZ* transgenic reporter analysis in mouse embryos at E11.5, a stage characterized by wide-spread and functionally relevant *Shox2* expression in multiple tissues^19,47^ (**Fig. 1B, C**). This analysis included the validation of 16 genomic elements (DE +329kb and +331kb were part of a single reporter construct) and revealed reproducible enhancer activities in 12/16 cases (**Figs. 1B, S1B and Table S2**). Most of the individual enhancer activities localized to either craniofacial, cranial nerve, mid-/hindbrain or limb subregions known to be dependent on *Shox2* expression and function^27,28,30,31^ (**Fig. 1C**). For example, DE9 (+475kb) and DE15 (+606kb), both exhibiting limb and craniofacial H3K27ac marks, drove LacZ reporter expression exclusively in *Shox2*-overlapping craniofacial domains in the medial nasal (MNP) and maxillary-mandibular (MXP, MDP) processes, respectively (**Fig. 1C)**. In line with DE15 activity, *Shox2* expression in the MXP-MDP junction is known to be required for temporomandibular joint (TMJ) formation in jaw morphogenesis^28^. DE1, 5 and 12 instead showed activities predominantly in cranial nerve tissue, including the trigeminal (TGn), facial (FGn) and jugular (JGn) ganglia, as well as the dorsal root ganglia (DRG) (**Fig. 1C**). *Shox2* is expressed in all these neural crest-derived tissues, but a functional requirement has only been demonstrated for FGn development and the mechanosensory neurons of the DRG^30,32^. While no H3K27ac profiles for cranial nerve populations were available from ENCODE^56^, both DE5 and DE12 elements showed increased H3K27ac in craniofacial compartments at E11.5 (**Fig. 1B**), likely reflecting the common neural-crest origin of a subset of these cell populations^57^. At mid-gestation, *Shox2* is also expressed in the diencephalon (DiE), midbrain (MB) and hindbrain (HB), and is specifically required for cerebellar development^31^. Gene desert enhancer assessment also identified a set of brain enhancers (DE7, 14 and 16) overlapping *Shox2* domains in the DiE, MB and/or HB. (**Fig. 1C**). Although H3K27ac marks were present in limbs at most predicted DEs, only two elements (DE6 and DE10) drove LacZ reporter expression in the E11.5 limb mesenchyme in a sub-regionally or limb type-restricted manner, respectively (**Fig. 1C**). However, despite elevated cardiac H3K27ac in a subset of DEs, none of the validated elements drove reproducible LacZ reporter expression in the heart at E11.5 (**Fig. 1B, 1C**). Taken together, our *in vivo* enhancer-reporter screen based on systematic epigenomic profiling and transgenic reporter validation identified multiple DE elements with *Shox2*-overlapping activities, pointing to a role of the gene desert as an enhancer hub directing pleiotropic *Shox2* transcription.

### The *Shox2* gene desert shapes a chromatin loop with tissue-specific features

Recent studies have shown that sub-TAD interactions can be pre-formed or dynamic, and that 3D chromatin topology can affect enhancer-promoter communication in distinct cell types or tissues^58–60^. To explore the 3D chromatin topology across the *Shox2* TAD and flanking regions, we performed region capture HiC (C-HiC) targeting a 3.5Mb interval in dissected E11.5 mouse embryonic forelimbs, mandibles, and hearts, tissues known to be affected by *Shox2* loss-of-function (**Figs. 2A, S2A**). C-HiC contact maps combined with analysis of insulation scores to infer inter-domain boundaries revealed a tissue-invariant *Shox2*-containing TAD that matched the extension observed in mESCs^61^ (**Figs. 2A, S2A, Table S3**). C-HiC profiles further showed sub-TAD organization into *Shox2*-flanking upstream (U-dom) and downstream (D-dom) domains as hallmarked by loop anchors and insulation scores, with the D-dom spanning almost the entire gene desert (**Figs. 2A, S2A**). Virtual 4C (v4C) using a viewpoint centered on the *Shox2* transcriptional start site (TSS) further demonstrated confinement of *Shox2*-interacting elements to U-dom and D-dom intervals, or TAD boundary regions (**Fig. 2A, Table S3**). Remarkably, the most distal D-dom compartment spanning ∼170kb revealed dense chromatin contacts restricted to heart tissue and delimited by weak insulation boundaries which co-localized with non-convergent CTCF sites (**Fig. 2A, B, S2A**). While this high-density contact domain (HCD) contained the majority of the previously identified (non-cardiac) gene desert enhancers (DE5-12), subtraction analysis further corroborated increased chromatin contacts across the HCD and domain insulation specifically in cardiac tissue as opposed to limb or mandibular tissue, indicating a potentially repressive function due to condensed chromatin state (**Fig. 2A-C, S2B**). However, no region-specific accumulation of repressive histone marks (H3K27me3 or H3K9me3) was observed in whole heart samples from ENCODE (**Fig. S3A, B**). Instead, V4C subtraction analysis with defined viewpoints on positively validated DEs indicated that specifically in heart tissue, enhancer elements outside the HCD (DE1, 15) were reduced in contacts with elements inside (**Fig. S3C**). In turn, enhancer viewpoints inside the HCD (DE5, 9, 10) showed reduced contacts with elements outside (**Fig. S3C**). Collectively, our results imply that *Shox2* is preferentially regulated by upstream (U-dom) and downstream (D-dom) regulatory domains that contain distinct sets of active tissue-specific enhancers. Hereby, the gene desert forms a topological chromatin environment (D-dom) that in specific tissue context might enable certain enhancers to interact more efficiently with the *Shox2* promoter.

**Figure 2.**
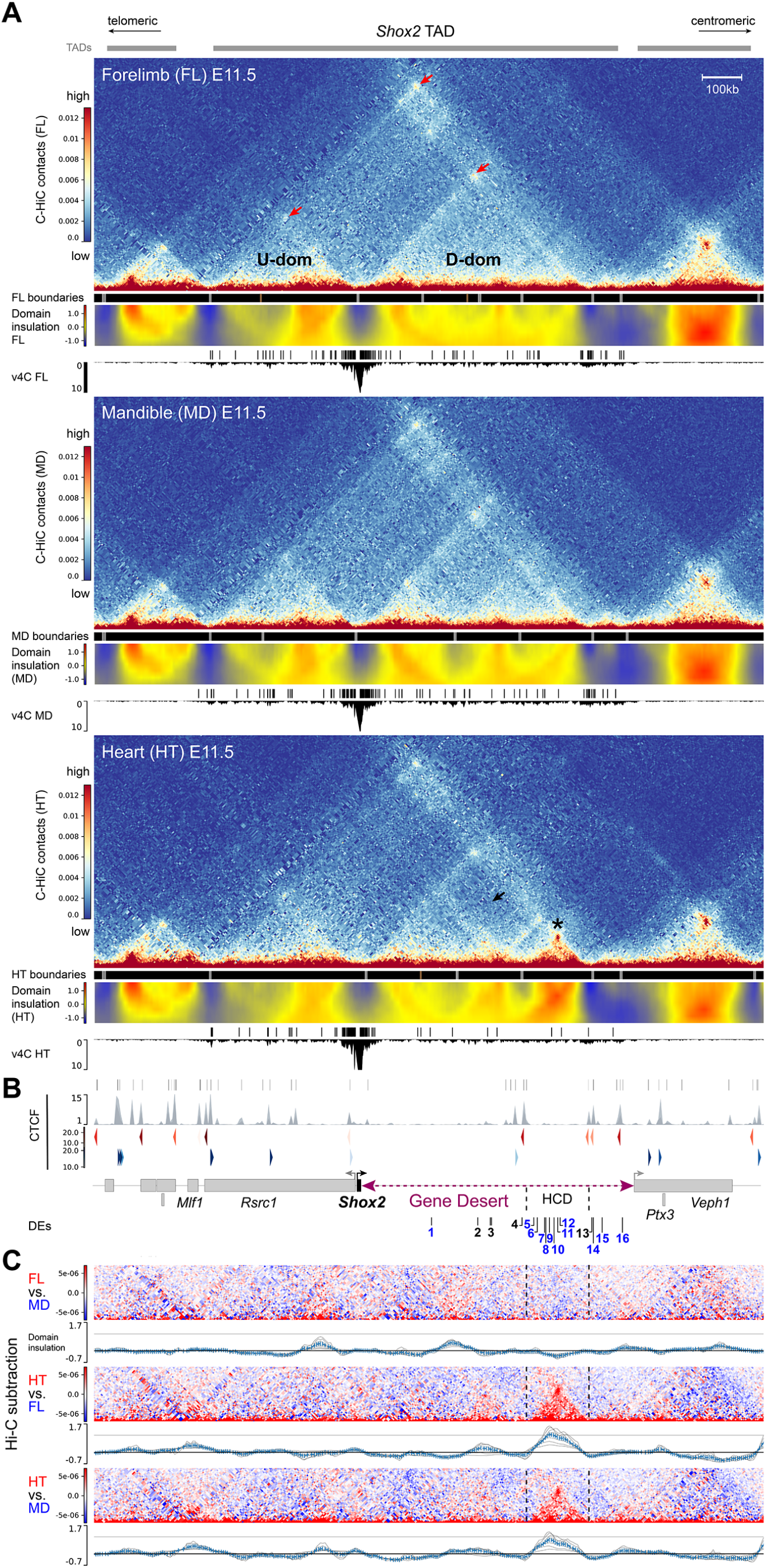
3D chromatin architecture across the *Shox2* regulatory landscape in distinct tissues. (**A**) C-HiC analysis of the genomic region containing the *Shox2* TAD^61^ in wildtype mouse embryonic forelimb (FL), mandible (MD) and heart (HT) at E11.5 (see also **Fig. S2**). The chr3:65977711-67631930 (mm10) interval is shown. Upper panels (for each tissue): Hi-C contact map revealing upstream (U-dom) and downstream (D-dom) domains flanking the *Shox2* gene. Middle panels: Stronger (gray boxes, p < 0.01) and weaker (brown boxes, p > 0.01, < 0.05) domain boundaries based on TAD separation score. A matrix showing normalized inter-domain insulation score (blue = weak insulation, red = strong insulation) is plotted below. Bottom panels: Virtual 4C (v4C) using a *Shox2*-centered viewpoint shows *Shox2* promoter interaction profiles in the different tissues. *Shox2* contacting regions (q < 0.1) as determined by GOTHiC^126^ are shown on top. Red arrows point to chromatin domain anchors. Asterisk marks a high-density contact domain (HCD) observed only in heart tissue (chr3:66402500-66572500). Black arrow indicates reduction of internal D-dom contacts between elements inside the HCD and outside in the heart sample (see also **Fig. S2**). (**B**) Top: CTCF enrichment in mESCs^61^. Bottom: CTCF motif orientation (red/blue) and strength (gradient). Protein coding genes (gene bodies) are indicated below. DEs, predicted gene desert enhancers validated in Fig. 1 (blue: tissue-specific activity). (**C**) C-HiC subtraction to visualize tissue-specific contacts for each tissue comparison (red/blue). Plots below display the corresponding subtracted inter-domain insulation scores. Dashed lines demarcate the HCD borders.

### Control of pleiotropic *Shox2* dosage and embryonic survival by the gene desert

To explore the functional relevance of the gene desert as an interactive hub for *Shox2* enhancers in mouse embryos, we used CRISPR/Cas9 in mouse zygotes to engineer an intra-TAD gene desert deletion allele (GD^Δ^) (**Figs. 3A, S4A, B; Tables S4, S5**). F1 mice heterozygous for this allele (GD^Δ/+^) were born at expected Mendelian ratios and showed no impaired viability and fertility. Following intercross of GD^Δ/+^ heterozygotes we compared *Shox2* transcripts in GD^Δ/Δ^ and wildtype (WT) control embryos, with a focus on tissues marked by DE activities (**Figs. 1C, 3B-E**). Despite loss of at least three enhancers with limb activities (hs1262, DE6, DE10), *Shox2* expression was still detected in fore- and hindlimbs of GD^Δ/Δ^ embryos, as determined by *in situ* hybridization (ISH) (**Fig. 3B**), albeit at ∼50% reduced levels as determined by RT-qPCR (**Fig. 3C, Table S6**). These results point to a functional role of the gene desert in ensuring robust *Shox2* dosage during proximal limb development^27^. *Shox2* expression in distinct craniofacial compartments was more severely affected by the loss of the gene desert (**Fig. 3D, E**). Downregulation of *Shox2* transcripts was evident in the MNP, anterior portion of the palatal shelves, and the proximal MXP-MDP domain of GD^Δ/Δ^ embryos at E10.5 and E11.5, compared to wild-type controls (**Fig. 3D**). Concordantly, and in contrast to *Rsrc1* mRNA levels which remained normal, *Shox2* was depleted in the nasal process (NP) and MDP of GD^Δ/Δ^ embryos at E11.5 (**Fig. 3E**). Strikingly, these affected subregions corresponded to the activity domains of the identified DE9 (MNP) and DE15 (MXP-MDP) gene desert enhancers indicating essential craniofacial *Shox2* regulation (**Figs. 1C, 3F**). Taken together, these results demonstrate a critical functional role of the gene desert in transcriptional regulation of *Shox2* during craniofacial and proximal limb morphogenesis^27,28,33^. While our transgenic analysis also uncovered DEs with activities in brain or cranial nerve regions (**Fig. 1C**), no overt reduction in spatial *Shox2* expression was detected in corresponding subregions in GD^Δ/Δ^ embryos (**Fig. 3D**). This is likely attributed to the presence of *Shox2*-associated brain enhancers located in the U-dom (e.g., hs1413) or downstream of the D-dom and the deleted gene desert interval (e.g., DE16) which show partially overlapping activities.

**Figure 3.**
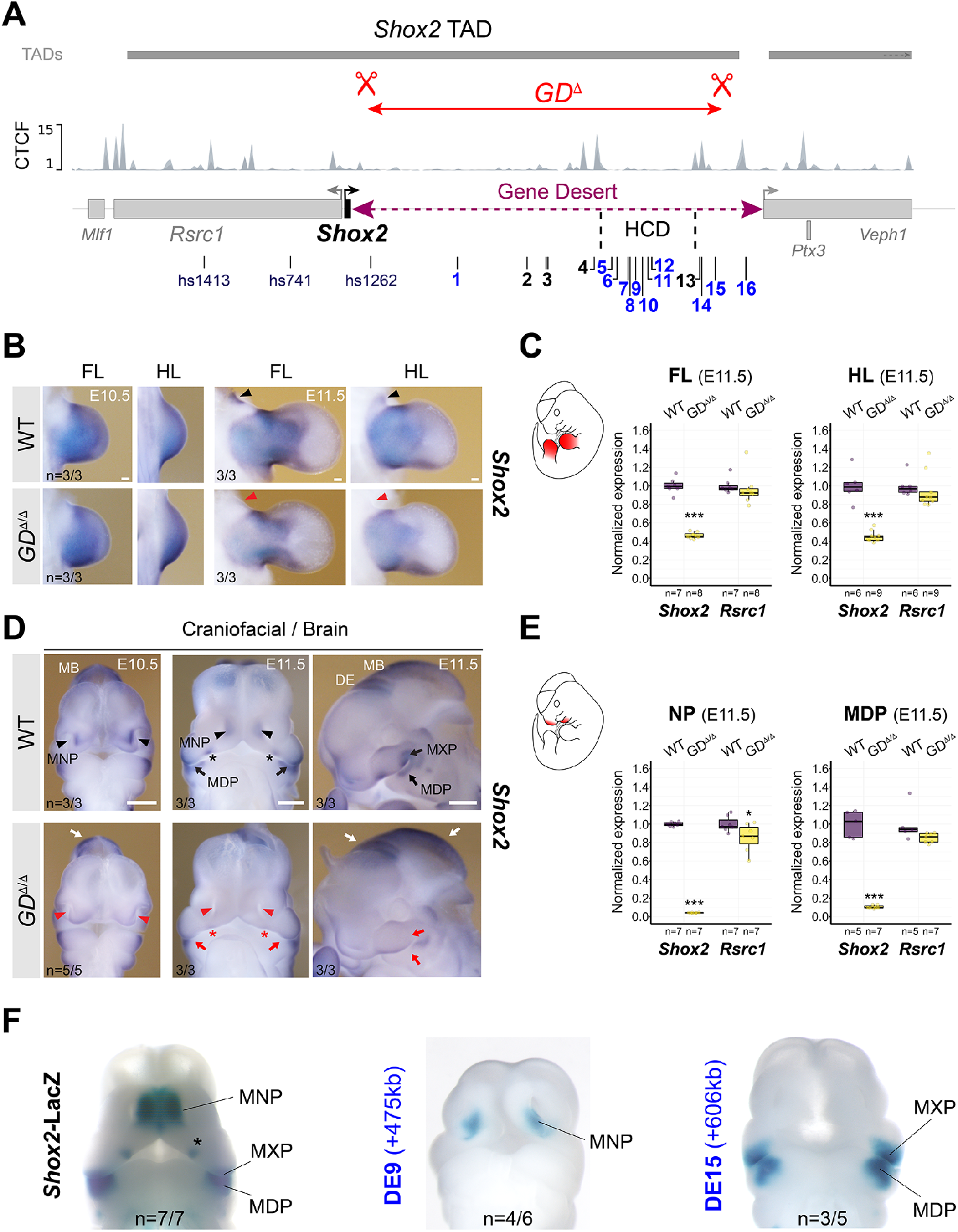
Gene desert deletion reduces *Shox2* in limb and craniofacial compartments. (**A**) CRISPR/Cas9-mediated deletion of the intra-TAD *Shox2* gene desert interval (GD^Δ^) (mm10, chr3:66365062-66947168). Vista (hs) and newly identified gene desert enhancers (1-16, active in blue) are displayed along with TAD interval and CTCF peaks from mESCs^61^. HCD, high-density contact domain (see Fig. 2). (**B**, **D**) ISH revealing spatial *Shox2* expression in fore- and hindlimb (FL/HL), craniofacial compartments, and brain in GD^Δ/Δ^ embryos compared to wildtype (WT) controls at E10.5 and E11.5. Red arrowheads and red arrows point to regions with severely downregulated or reduced *Shox2* expression, respectively. Red asterisk demarcates *Shox2* loss in the anterior portion of the palatal shelves. White arrows indicate regions (diencephalon, DE and midbrain, MB) without overt changes in *Shox2* expression. Scale bars, 500 gm (B) and 100 gm (D). (**C, E**) Quantitative mRNA analysis (qPCR) in limb and craniofacial tissues of WT and GD^Δ/Δ^ embryos. Box plots indicate interquartile range, median, maximum/minimum values (bars). Dots represent individual data points. ***, P < 0.001; *, P < 0.05 (two-tailed, unpaired t-test for qPCR). (**F**) DE9 and DE15 enhancer activities (Fig. 1C) overlap *Shox2* expression in medial nasal process (MNP) and maxillary-mandibular (MXP-MDP) regions, respectively, in mouse embryos at E11.5. Asterisk marks anterior palatal shelf. “n” indicates number of embryos per genotype or transgene analyzed, with similar results.

Despite the lack of identification of any *in vivo* heart enhancers in the gene desert following transgenic reporter analysis from epigenomic whole-heart predictions (**Fig. 1C**), spatial and quantitative mRNA analysis in GD^Δ/Δ^ embryos revealed absence of *Shox2* transcripts from the cardiac sinus venosus (SV) that harbors the population of SAN pacemaker progenitors^62^ (**Fig. 4A, B**). In accordance with the essential role of *Shox2* in the differentiation of SAN progenitors and the related lethality pattern in *Shox2*-deficient mouse embryos^34^, cardiac *Shox2* depletion in GD^Δ/Δ^ embryos triggered arrested development and embryonic lethality at around E12 (n=5/5) (**Fig. S4C**). Immunofluorescence further confirmed lack of Shox2 protein in the Hcn4-positive domain of SAN pacemaker cells in the SV of GD^Δ/Δ^ hearts compared to WT controls at E11.5 (**Fig. 4C, S4D**). Together, these results demonstrated a requirement of the gene desert for embryonic viability directly associated with transcriptional control of cardiac *Shox2*.

**Figure 4.**
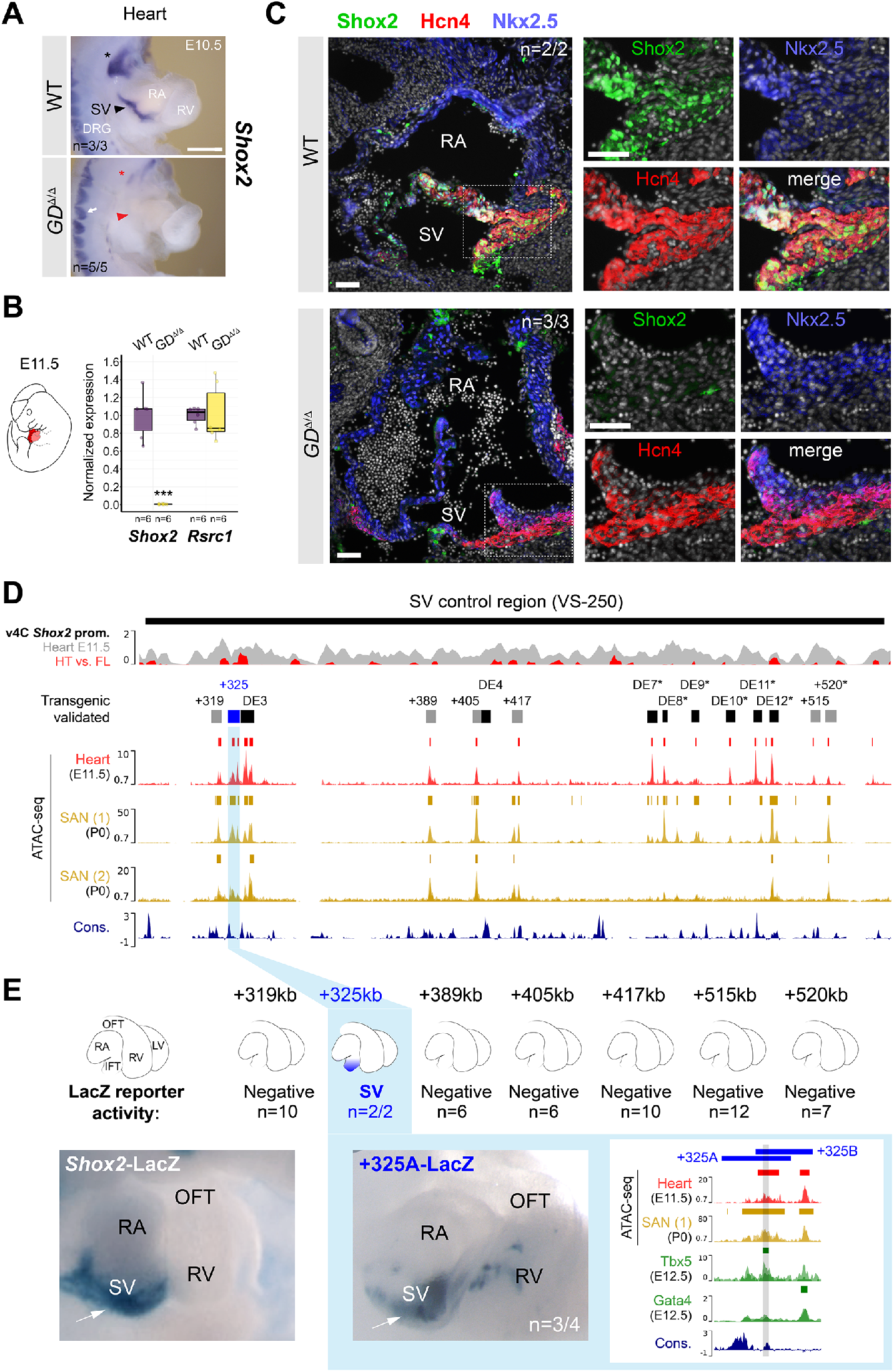
The gene desert controls cardiac *Shox2* essential for embryonic viability. (**A**) ISH revealing absence of *Shox2* transcripts in the cardiac sinus venosus (SV) of GD^Δ/Δ^ embryos at E10.5 (red arrowhead). Red asterisk points to reduced *Shox2* in the nodose ganglion of the vagus nerve. White arrow indicates normal *Shox2* expression in the dorsal root ganglia (DRG) of GD^Δ/Δ^ embryos. Scale bar, 100 pm. (**B**) Quantitative PCR (qPCR) revealing depletion of *Shox2* in GD^Δ/Δ^ hearts compared to WT controls at E11.5. Box plots indicate interquartile range, median, maximum/minimum values (bars). Dots represent individual data points. ***P < 0.001 (two-tailed, unpaired t-test). (**C**) Co-localization of Shox2 (green), Hcn4 (red) and Nkx2-5 (blue) in hearts of GD^Δ/Δ^ and WT control embryos at E11.5. Shox2 is lost in the Hcn4-marked SAN pacemaker myocardium in absence of the gene desert (dashed outline). Nuclei are shown in gray. Scale bars, 50p.m. **(D)** SAN enhancer candidate regions in the gene desert interval (VS-250) essential for *Shox2* expression in the SV^63^. Top: Virtual 4C (v4C) *Shox2* promoter interaction signature in embryonic hearts (gray) overlapped with heart (HT) versus forelimb (FL) subtraction profiles (red) (*see also* **Fig. S3A**). Below: ATAC-seq profiles and peak-calls from embryonic hearts at E11.5 (this study) and SAN pacemaker cells from sorted *Hcn4*-GFP mouse hearts at P0 (Fernandez-Perez *et al*. (1)^64^ and Galang *et al*. (2)^44^). Desert enhancers (DEs) accessible in SAN progenitors and additional predicted SAN enhancer elements are shown as black and gray bars, respectively. Distance to the *Shox2* TSS in kb (+) is indicated. Cons, vertebrate conservation track by PhyloP. (**E**) LacZ transgenesis identifies a 3.9kb genomic element 325kb downstream of the *Shox2* TSS (+325) able to drive LacZ reporter expression in the SV at E11.5 (**Fig. S5**). 325A and 325B subregions each drive *Shox2* overlapping SV activity (see **Fig. S5** for 325B). The conserved interval in the 325A/B overlapping region shows a Tbx5 peak (gray bar) but no significant Gata4 enrichment in embryonic hearts at E12.5^65,66^ (green tracks). “n” denotes fraction of biological replicates with reproducible results. Single numbers represent total number of transgenic embryos analyzed (without reproducible staining in the heart). RA, right atrium. RV, right ventricle. OFT, outflow tract.

### Resilient gene desert enhancer architecture ensures robust cardiac *Shox2* expression

Abrogation of *Shox2* mRNA in the SV of GD^Δ/Δ^ embryos implied the presence of enhancers with cardiac activities, similar to the regulation of other TFs implicated in the differentiation of SAN progenitor cells^44^. In agreement with our findings, a recent study^63^ has reported that deletion of a 241kb interval within the gene desert (VS-250, mm10 chr3:66444310-66685547) is sufficient to deplete *Shox2* in the SV. This resulted in a hypoplastic SAN and abnormally developed venous valve primordia responsible for embryonic lethality^63^. We therefore concluded that loss of *Shox2* in hearts of GD^Δ/Δ^ embryos results from inactivation of one (or more) SV/SAN enhancer(s) in the VS-250 interval (**Fig. 4D**). While our epigenomic analysis from whole hearts identified multiple elements with heart enhancer signatures (H3K27ac) (**Fig. 1B**), none was found to drive reproducible cardiac activity in embryos at E11.5 by transgenic reporter analysis (**Fig. 1C**). To refine *Shox2*-associated cardiac enhancer predictions we performed ATAC-seq from mouse embryonic hearts at E11.5 and intersected the results with reprocessed open chromatin signatures from HCN4^+^-GFP sorted SAN pacemaker cells of mouse hearts at P0, available from two recent studies^44,64^ (**Fig. 4D**, **Table S7**). Intersection of peak calls within the VS-250 interval identified multiple sites with overlapping accessible chromatin in embryonic hearts and perinatal SAN cells. While a subset of these candidate SAN enhancer elements overlapped DEs validated for non-cardiac activities (DE 3, 4, 7-12), the remaining ATAC-called elements (+319, +325, +389, +405, +417, +520) included yet uncharacterized elements showing variable enrichment for Tbx5, Gata4 and/or Tead TFs which are associated with SAN enhancer activation^44,65,66^ (**Figs. 4D, S5A, Table S7**). To obtain complete functional validation coverage, we subjected these new putative SV/SAN enhancer elements to LacZ reporter transgenesis in mouse embryos (**Fig. 4D, Table S8**). This analysis identified a single element located 325kb downstream of *Shox2* (+325) that was able to drive reproducible LacZ reporter expression in the cardiac SV in a reproducible manner (**Fig. 4D, S5B**). To further define the core region responsible for the SV-specific activity we divided the 4kb-spanning +325 module into two elements: +325-A and +325-B (**Fig. 4E, Table S8**). These elements overlapped in a conserved block of sequence (1.5kb) that showed an open chromatin peak in embryonic hearts at E11.5 and SAN cells at P0, and also co-localized with Tbx5 enrichment at E12.5 (**Fig. 4E**). Both +325-A and +325-B elements retained SV enhancer activity on their own in transgenic reporter assays, indicating that the core sequence is responsible for SV activity (**Fig. 4E, S5B**). To identify cardiac TF interaction partners in enhancers at the motif level, we then established a general framework based on a former model of statistically significant matching motifs^67^ and restricted to TFs expressed in the developing heart at E11.5 (**Table S9**) (see ***Methods***). This approach identified a bi-directional Tbx5 motif in the active core [*P*=1.69e-05 (+) and *P*=1.04e-05 (-)] of the +325 SV enhancer module which in addition with ChIP-seq binding suggested direct interaction of Tbx5 (**Fig. 4E, S5C**). In contrast, no motifs or binding of other established cardiac *Shox2* upstream regulators (e.g., Isl1) were identified in this core sequence (**Fig. S5A, S5C**). In summary, our results identified the +325 module as a remote Tbx5-interacting cardiac enhancer associated with transcriptional control of *Shox2* in the SV and thus likely required for SAN progenitor differentiation^34,63^.

The mouse +325 SV enhancer core module is conserved in the human genome where it is located 268kb downstream (+268) of the TSS of the *SHOX2* orthologue. Taking advantage of fetal left and right atrial (LA and RA) as well as left and right ventricular (LV and RV) tissue samples at post conception week 17 (pcw17) available from the Human Developmental Biology Resource at Newcastle University, we conducted H3K27ac ChIP-seq and RNA-seq to explore chamber-specific SV enhancer activity during pre-natal human heart development (**Fig. 5A**). These experiments uncovered an atrial-specific H3K27ac signature at the (+268) conserved enhancer module, matching the transcriptional specificity of *SHOX2* distinct from the ubiquitous profile of *RSRC1* in human hearts (**Fig. 5A**). This result indicating human-conserved activity prompted us to investigate the developmental requirement of the SV enhancer *in vivo*. Therefore, we used CRISPR-Cas9 in mouse zygotes (CRISPR-EZ)^68^ to delete a 4.4kb region encompassing the +325 SV enhancer interval (SV-Enh^Δ^) (**Figs. 5B, S5D, E; Tables S4, S5**). F1 mice heterozygous for the SV enhancer deletion (SV-Enh^Δ/+^) were phenotypically normal and subsequently intercrossed to produce homozygous SV-Enh^Δ/Δ^ embryos. ISH analysis pointed to downregulation of *Shox2* transcripts in the SV region in SV-Enh^Δ/Δ^ embryos at E10.5 and qPCR analysis at the same stage demonstrated a ∼60% reduction of *Shox2* in hearts of SV-Enh^Δ/Δ^ embryos compared to WT controls (**Fig. 5C**). Despite this reduction of *Shox2* dosage in embryos, SV-Enh^Δ/Δ^ mice were born at normal Mendelian frequency and showed no overt phenotypic abnormalities during adulthood. Together, these results imply that multiple gene desert enhancers are in control of *Shox2* expression in SAN progenitors and establish that in such a system the +325 SV enhancer acts as a core module required for buffering of cardiac *Shox2* to protect from dosage-reducing mutations.

**Figure 5.**
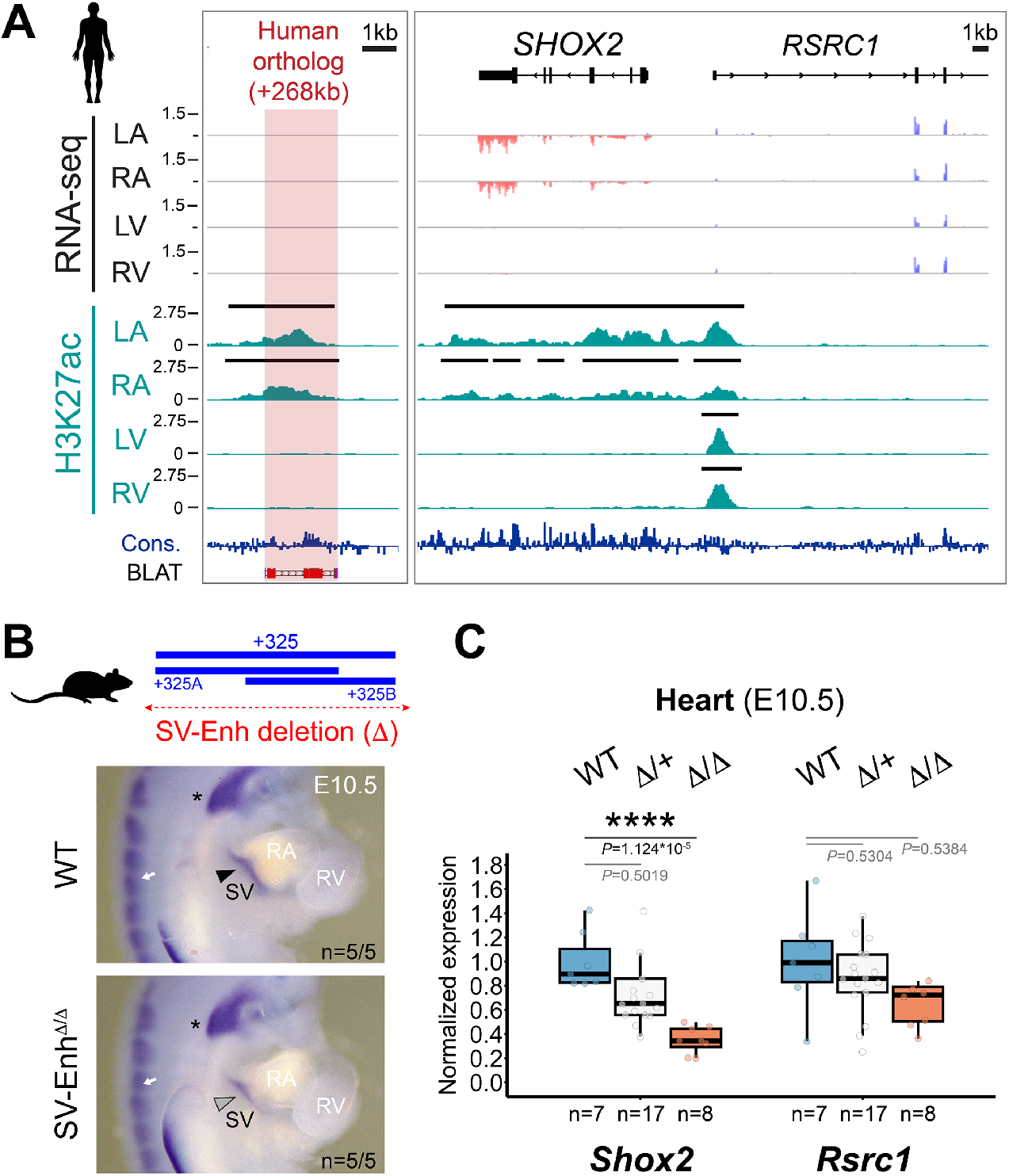
Enhancer-mediated transcriptional robustness safeguards *Shox2* in the heart. (**A**) H3K27 acetylation ChIP-seq (H3K27ac) and RNA-seq profiles from human fetal heart compartments at PWC17 across the human orthologous sequence of the +325-mouse sinus venosus (SV) enhancer and the *SHOX2* interval. The left ventricle (LV) dataset has been previously published^143^. +268, distance to *SHOX2* TSS. Cons, mammalian conservation by PhyloP. (**B**) Top: Generation of a +325 SV enhancer deletion (4.4kb) allele in mice (SV-Enh^Δ^). Below: *Shox2* mRNA distribution (ISH) in SV-Enh^Δ/Δ^ compared to WT mouse embryos at E10.5. (**C**) qPCR analysis of *Shox2* and *Rsrcl* mRNA levels in SV-Enh^Δ/+^ and SV-Enh^Δ/Δ^ embryonic hearts at E10.5 compared to WT controls. Box plot indicates interquartile range, median, maximum/minimum values (bars) and individual biological replicates (n). *P*-values are shown, with **** P<0.0001 (two-tailed, unpaired t-test). Three outliers, two datapoints of *Shox2* A/+ replicates and one for *Rsrcl* (A/A), are outside of the scale shown. “n” indicates number of biological replicates analyzed, with similar results. LA, left atrium. RA, right atrium. RV, right ventricle.

### A gene desert limb enhancer repertoire promotes stylopod morphogenesis

Given another essential role of *Shox2* in proximal limb development we next addressed the phenotypic requirement of the gene desert for skeletal limb morphogenesis. *Shox2* is essential for stylopod formation and thus analysis of skeletal elements serves as an ideal readout for the study of enhancer-related *Shox2* dosage reduction in the proximal limb^19,27^. Neither knockout of the hs1262 proximal limb enhancer^19^ nor the identification of new limb enhancers (DE6, DE10) located in the gene desert (**Fig. 1**) was sufficient to explain the ∼50% *Shox2* reduction observed in proximal fore-(FL) and hindlimbs (HL) of GD^Δ/Δ^ embryos (**Fig. 3B, C**). To refine our epigenomic limb enhancer predictions at the spatial level we reprocessed previously published ChIP-seq datasets from dissected proximal and distal limbs at E12^55^ which revealed multiple proximal-specific H3K27ac peaks (**Fig. 6A**). These included several elements not significantly enriched in H3K27ac maps from whole-mount limb tissue (**Fig. 1B**). Interestingly, multiple elements marked by H3K27ac in proximal limbs also showed H3K27me3 in distal limb mesenchyme reflecting compartment-specific bivalent epigenetic regulation^55^. With the goal to identify the complement of H3K27ac-marked elements that interact with the *Shox2* promoter we next performed circular chromosome conformation capture (4C-seq) with a *Shox2* viewpoint from dissected proximal limbs at E12.5 (**Fig. 6B, Table S10**). Processing of two replicates resulted in reproducible peaks which confirmed physical interaction between the *Shox2* promoter and each of the *bona-fide* proximal limb enhancers characterized previously: hs741 located in the upstream domain (U-dom) and hs1262 located in the gene desert (D-dom)^19,47^ (**Fig. 6B, C**). Other prominent 4C-seq peaks in the gene desert co-localized with either previously validated enhancer elements with non-limb activities at E11.5 (DE1, 4, 6, 9, 15) or non-validated elements with proximal limb-specific H3K27ac enrichment (+237kb and +568kb) (**Fig. 6B, C**). Open chromatin and H3K27ac profiles further indicated that the *Shox2*-interacting DE4 (+407) element was unique in its H3K27ac pattern (initiated at E11.5), while other proximal limb (candidate) enhancers showed activity marks already at E10.5. Therefore, we decided to analyze the spatiotemporal enhancer dynamics of newly identified (+237kb, +568kb) and temporally dynamic (+407) limb candidate enhancer regions using stable transgenic LacZ reporter mouse lines. For comparison, we also assessed the previously validated hs741 and hs1262 *Shox2* limb enhancers^19,47^ (**Fig. 6C, S6A and Table S11**). Remarkably, at E12.5, each element on its own was able to drive reporter expression in the proximal fore- and hindlimb mesenchyme in a pattern overlapping *Shox2*, establishing a complement of at least five proximal limb enhancers (PLEs) that contact *Shox2*, four of which reside within the gene desert (PLE2-PLE5) (**Fig. 6C, S6A**). These activity patterns generally showed strong reporter signal in the peripheral mesenchyme of the stylopod and zeugopod elements (**Fig. 6C, S6A**). *Shox2* expression is progressively downregulated within the differentiating chondrocytes of the proximal skeletal condensations of the limbs from E11.5, while its expression remains high in the surrounding mesenchyme and perichondrium^49,69–71^. In accordance, activities of the newly discovered elements (PLE3-5) remained excluded from the chondrogenic cores of the skeletal condensations, consistent with a role in generating the *Shox2* expression pattern required for stylopodial chondrocyte maturation and subsequent osteogenesis^11,27^. PLE3 (+237) was initiated in the proximal limb mesenchyme at E11.5 with persistent signal until E13.5 and most closely recapitulating the late *Shox2* expression pattern^27,49^ (**Fig. S6A**). Instead, PLE4 (+407) drove reporter activity already at E10.5 in a more widespread pattern leaking into distal forelimbs at later stages, in line with H3K27ac enrichment in distal forelimbs (**Figs. 6A, S6A**). PLE5 (+568) was initiated only at E12.5 and its activity remained restricted to the proximal-anterior (**Fig. S6A**). Together, these diverse and partially overlapping enhancer activities pointed to dynamic interaction of *Shox2* gene desert enhancers during limb development. In addition, to achieve insight into PLE configuration at the chromatin level we performed 4C-seq with viewpoints at PLE2 and PLE4 which indicated the formation of a complex involving PLE1, 3 and 4, but not PLE2 (**Fig. S6B-D**). These findings indicate that PLE interactions might not necessarily be restricted to U-dom or D-dom sub-compartments for *Shox2* regulation in the limb. Taken together, these results identify the gene desert as a multipartite *Shox2* limb enhancer unit and indicate an instructive role in the transcriptional control of stylopod morphogenesis.

**Figure 6.**
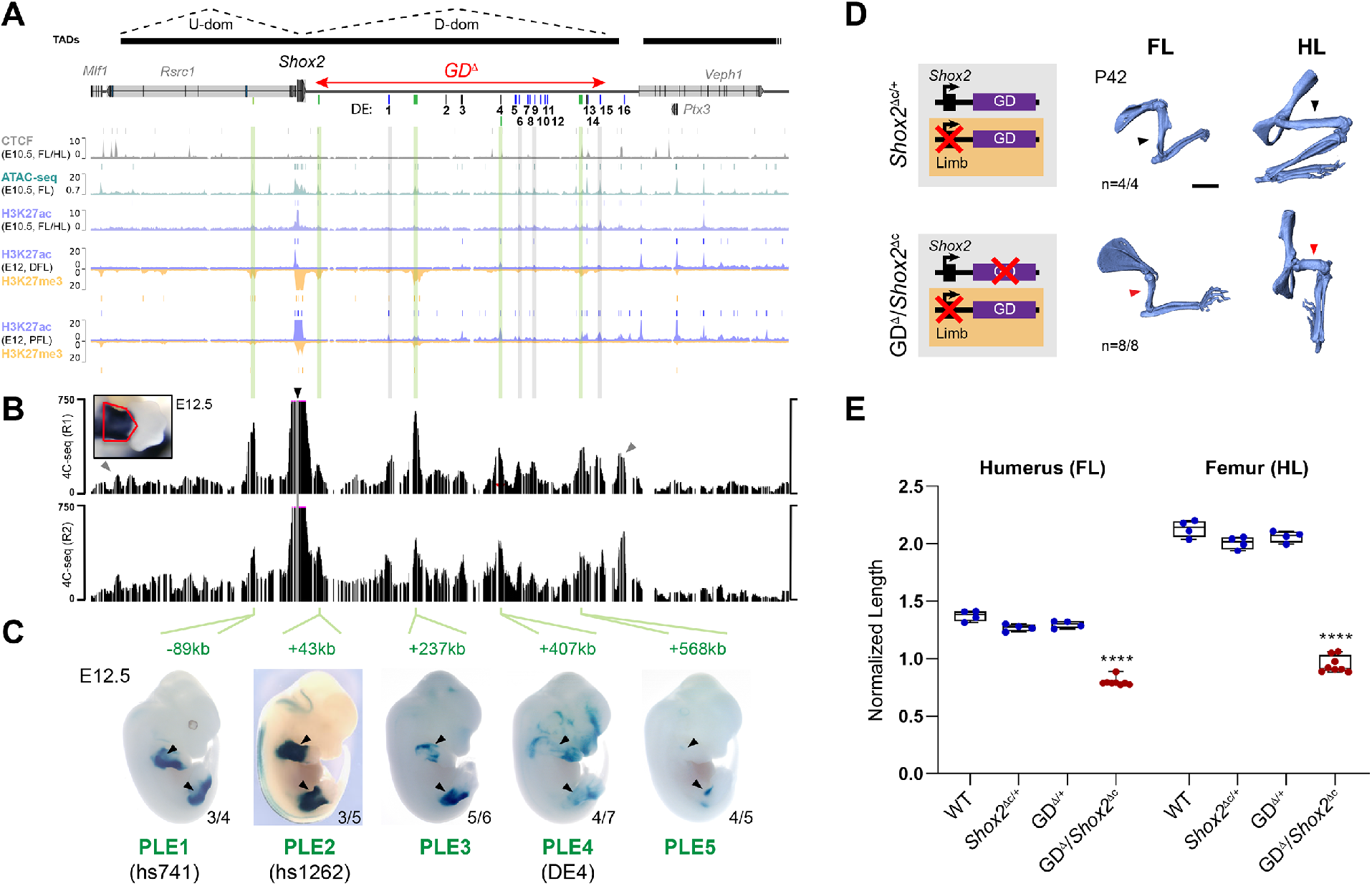
Stylopod morphogenesis is dependent on distributed proximal limb enhancers in the gene desert. (**A**) Re-processed ChIP-seq datasets from mouse embryonic limbs at E10.5 (CTCF, ATAC-seq, H3K27ac) and E12 (H3K27ac, H3K27me3) showing epigenomic profiles at the *Shox2* locus^55,60,130^. Bars above each track represent peak calls across replicates. DFL, distal forelimb. PFL, proximal forelimb. On top: TAD extension in mESCs^61^ (black bars) with desert enhancers identified in Fig. 1 (DEs 1-16; blue indicates validated activity). Red double arrow demarcates the deleted gene desert interval (GD^Δ^). (**B**) 4C-seq interaction profiles from two independent biological replicates (R1, R2) of proximal limbs at E12.5 (red outline). Black arrow indicates the 4C-seq viewpoint at the *Shox2* promoter. Gray arrowheads point to CTCF-boundaries of the *Shox2*-TAD. Green lines (in **A**) indicate *Shox2*-interacting elements with putative proximal limb activities. Gray lines mark 4C-seq peaks overlapping previously validated DEs without such activities (Fig. 1C). (**C**) Identification of proximal limb enhancers (PLEs) through transgenic LacZ reporter assays in mouse embryos at E12.5. Embryos shown are representatives from stable transgenic LacZ reporter lines (**Fig. S6A**). Reproducibility numbers from original transgenic founders are listed for each element (bottom right). (**D**) Left: Schematics illustrating gene desert inactivation (GD^Δ^) in the presence of reduced limb *Shox2* dosage based on *Prxl*-Cre-mediated *Shox2* deletion (*Shox2*^Δc^). Right: Micro-CT scans of fore (FL)- and hindlimb (HL) skeletons of GD^Δ^/*Shox2*^Δc^ and *Shox2*^Δc/+^ control mice at postnatal day 42 (P42). Red arrowheads point to severely reduced stylopods in GD^Δ^/*Shox2*^Δc^ individuals compared to controls (black arrowheads). “n”, number of biological replicates with reproducible results. (**E**) Micro-CT stylopod quantification at P42 reveals humerus and femur length reductions in GD^Δ^/*Shox2*^Δc^ mice. ****, P < 0.0001 (ANOVA).

To evaluate the functional and phenotypic contribution of the gene desert to stylopod formation we combined our gene desert deletion allele with a *Prx1*-Cre conditional approach for *Shox2* inactivation^27,72^. This enabled limb-specific conditional *Shox2* inactivation on one allele (*Shox2*^Δc^) paired with gene desert deletion on the other allele (GD^Δ^), allowing to bypass embryonic lethality caused by loss of cardiac *Shox2* (**Figs. 6D, S4**). Remarkably, this abolishment of gene desert-mediated *Shox2* regulation in limbs led to severe shortening of the stylopod with an approximate 60% reduction in humerus length and 80% decrease in femur extension in GD^Δ^/*Shox2*^Δc^ newborn mice (**Fig. S7A, B**). These skeletal abnormalities were in line with the stylopod phenotypes obtained by limb-specific *Shox2* dosage reductions in previous studies^19,27^. Concordantly, micro-computed tomography (pCT) from adult mouse limbs at P42 showed significant humerus length reduction of approximately 40% and decreased femur length of about 50% (**Fig. 6E**). Our results thus demonstrate an essential role of the gene desert in proximal limb morphogenesis and imply a significant functional contribution of the PLE2-5 modules to spatiotemporal control of *Shox2* dosage in the limb.

In summary, our study identifies the *Shox2* gene desert as an essential and dynamic chromatin unit that encodes an array of distributed tissue-specific enhancers that coordinately regulate stylopod formation, craniofacial patterning, and SAN pacemaker dependent embryonic progression (**Fig. 7A-C**). The arrangement of the enhancers appears modular but distributed in terms of tissue-specificities (**Fig. 7A**). While craniofacial and neuronal gene desert enhancers are hallmarked by driving mostly distinct subregional activities, limb enhancers (PLEs) show more overlapping activity domains, pointing to potential redundant intra-gene desert interactions. Hereby, the detection of a high-density contact domain (HCD) suggests that sub-TAD compartmentalization could further contribute to modulation of subregional enhancer activities (**Fig. 7B**). Finally, the demonstrated phenotypic requirement of the *Shox2* gene desert for multiple developmental processes underscores the importance of functional studies focused on the non-coding genome for better mechanistic understanding of congenital abnormalities (**Fig. 7C**).

**Figure 7.**
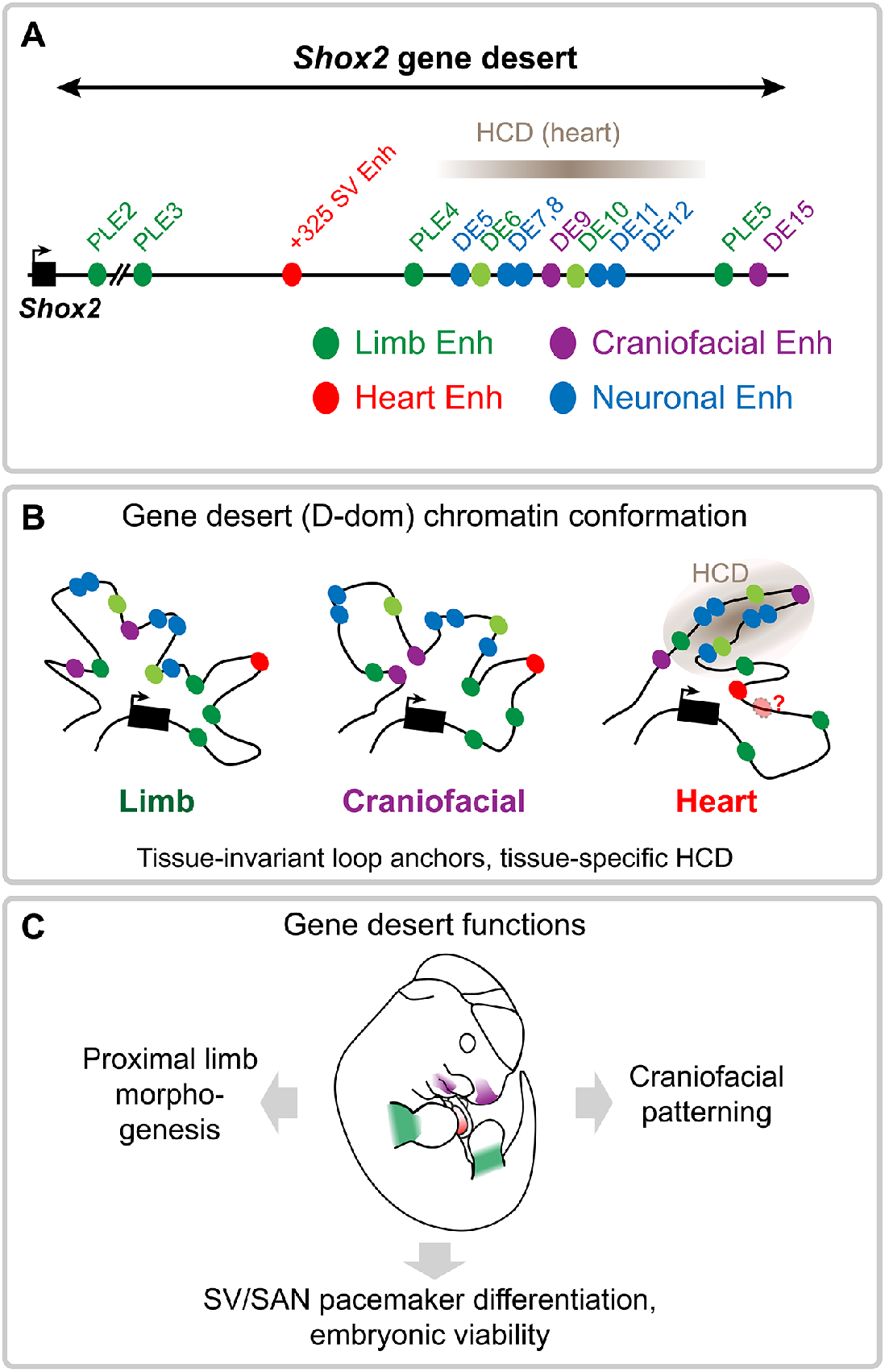
Graphical summary. (**A**) Identification of the *Shox2*-flanking gene desert as a reservoir for distributed transcriptional enhancers with activities in limb, craniofacial, cardiac, and neuronal cell populations. (**B**) The *Shox2* gene desert encodes distributed tissue-specific enhancers that are englobed in a dynamic chromatin domain (D-dom) with tissue-invariant loop anchors and a cardiac-specific high-density contact domain (HCD) that may influence enhancer activities. Additional gene desert enhancers are likely to participate in the regulation of cardiac *Shox2* in SAN progenitors. (**C**) Cumulative functions of gene desert enhancers orchestrate pleiotropic *Shox2* expression essential for proximal limb morphogenesis, craniofacial patterning, and cardiac pacemaker development.

## DISCUSSION

There is now evidence that dismantling of duplicates of ancient genomic regulatory blocks (GRBs) led to the emergence of gene deserts enriched in the neighborhood of regulatory genes such as TFs^73^. Functional assessment of TF gene deserts, including those in the *Hoxd* and *Sox9* loci, revealed that distal long-range enhancers represent critical *cis*-regulatory modules that control subregional expression domains through interaction with target gene promoters in a spatiotemporal manner^6,7,74,75^. Gene deserts can thus be conceived as genomic units coordinating dynamic enhancer activities in specific developmental processes, such as *HoxD*-dependent digit formation, and can be also hi-jacked by evolutionary processes to enable phenotypic diversification^9,11,76^. Silencer modules, insulating TAD boundaries and tethering elements (promoting long-range interactions) are involved in restriction or modulation of E-P interactions in metazoan genomes and can further contribute to gene desert functionality^77–79^. Recent studies also indicated that functional RNAs, such as IncRNAs or circRNAs, represent elements with enhancer-modifying or distinct regulatory potential within gene deserts^80^. Importantly, human disease-associated nucleotide variants in gene deserts are frequently linked to enhancer function, contributing to the spectrum of enhanceropathies^81–83^. Furthermore, deletions, inversions and duplications can alter or re-distribute interaction of gene desert enhancers with target gene promoters leading to congenital malformation or syndromes^10,17,18^. Despite these critical implications, the enhancer landscapes and related chromatin topology of most gene deserts near developmental genes remain incompletely characterized at the functional level^84^. In the current study, we addressed the functional necessity and *cis*-regulatory architecture of a gene desert flanking the *Shox2* transcriptional regulator, a critical determinant of embryogenesis and essential for limb, craniofacial and SAN pacemaker morphogenesis^37,47,85^. We identify the *Shox2* gene desert as a reservoir for highly subregional, tissue-specific enhancers underlying pleiotropic *Shox2* dosage by demonstrating essential contributions to stylopod morphogenesis, craniofacial patterning, and SV/SAN development. Our findings support a model in which gene deserts provide a scaffold for preferential chromatin domains that generate enhancer-mediated cell type or tissue-specific *cis*-regulatory output based on the integration of upstream signals.

Interpretation of gene desert function is dependent on accurate functional predictions of enhancer activities embedded in the genomic interval. Our approach using ChromHMM-filtered H3K27ac signatures from bulk tissues across a large range of embryonic stages (derived from ENCODE) serves as a baseline for the mapping of tissue-specific enhancer activities. However, while H3K27ac is known as the most specific canonical mark for active enhancers, it appears to not include all enhancers^86–88^. For example, recent studies evaluating H3K27ac-based tissue-specific enhancer predictions in mouse embryos revealed a substantial number of false-positives^54,89^. In turn, a significant fraction (∼14%) of validated *in vivo* enhancers were lacking enrichment of any canonical enhancer marks (ATAC-seq, H3K4me1, H3K27ac)^89^. In line with these observations, our transgenic reporter validation in many cases revealed more restricted or even distinct *in vivo* enhancer activities than those predicted by epigenomic marks. Such discrepancies might be partially originating from the use of bulk tissues or limited sensitivity of profiling techniques. In accordance, refinement of enhancer predictions using region-specific open chromatin data in combination with chromatin conformation capture (C-HiC, 4C-seq) enabled us to identify critical subregional cardiac and proximal limb enhancers missed by the initial epigenomic prediction approach.

Genomic deletion analysis uncovered an important functional role of the gene desert for pleiotropic expression and progression of embryonic development, the latter through direct control of *Shox2* in SAN pacemaker progenitors. Consistent with our findings, a parallel study narrowed the region essentially required for cardiac *Shox2* expression to a 241kb gene desert interval (termed VS-250)^63^. Here, we have identified a human-conserved SV enhancer (+325) located within this essential interval and specifically active in the SV/SAN region to maintain robust cardiac *Shox2* levels. These results add to recent progress in uncovering SAN enhancers of cardiac pacemaker regulators, including also *Isl1* or *Tbx3*^44,63^. Such findings not only shed light on the wiring of the GRNs driving mammalian conduction system development but also offer the opportunity to identify mutational targets linked to defects in the pacemaker system, such as arrhythmias^62^. Interestingly, removal of the *Isl1* SAN enhancer (ISE) in mice, as for our +325 *Shox2* enhancer, led to reduced target gene dosage but without subsequent embryonic or perinatal lethality^44^. These instances indicate that the GRNs orchestrating SAN pacemaker development are buffered at the *cis-*regulatory level, which can be enabled via partially redundant enhancer landscapes^19,90^. Similar to the binding profile of the +325 *Shox2* SV enhancer, a TF network involving Gata4, Tbx5 and Tead has been implicated in ISE activation, confirming a key role of Tbx5 in the activation of SAN enhancers in working atrial myocardium, while pacemaker-restricted identity may be established by repressive mechanisms^43,44,62^. ISE activity was also correlated with abnormal SAN function in adult mice and found to co-localize with resting heart rate SNPs, indicating potentially more sensitive GRN architecture in humans^44^. Intriguingly, coding and non-coding variants in the human *SHOX2* locus were recently associated with SAN dysfunction and atrial fibrillation, underscoring the value of human-conserved SAN enhancer characterization for functional disease variant screening^38,40,91,92^.

Arrangements of distributed enhancer landscapes conferring robust and cell type-specific transcription emerged as a common feature of metazoan gene regulatory architecture^93–95^. Gene deserts may thus not only function to promote robust expression boundaries and/or phenotypic resilience, but also represent a platform enabling evolutionary plasticity^9,73^. The conventional model of enhancer additivity based on individual small and stable regulatory contributions is likely predominant in gene deserts^96^. In support, we uncovered at least four *Shox2*-associated gene desert enhancers (PLE2-5) with overlapping activities in the proximal limb mesenchyme. Such regulatory architecture resembles the multipartite enhancer landscapes in *Indian Hedgehog* (*Ihh*) or *Gremlin1* loci, which as *Shox2* are involved in spatiotemporal coordination of proximal-distal limb identities with chondrogenic cues^22,24^. Our study further reveals gene desert enhancers with seemingly unique tissue specificities, such as the craniofacial DE9 and DE15 elements driving *Shox2*-overlapping reporter expression in the nasal process and maxillary-mandibular region, respectively. DE15 may be involved in jaw formation as *Shox2* inactivation in cranial neural crest cells in the maxilla-mandibular junction leads to dysplasia and ankylosis of the TMJ in mice^28^.

Our C-HiC experiments indicated that the repertoire of *Shox2* interacting elements (e.g., enhancers) is confined to the overarching TAD, without apparent cross-TAD boundary interactions^97^. The observed U-dom and D-dom assemblies (as evidenced by loop anchors) might reflect dynamic loop structures to facilitate *Shox2* promoter scanning similar to the organization at *HoxA* and *HoxD* loci that promotes nested and collinear gene expression^7,98^. C-HiC analysis also uncovered a high-density contact domain (HCD) emerging only in heart tissue. The absence of convergent CTCF sites flanking the HCD might reflect that a subset of contact domains form independently of cohesin-mediated loop extrusion, for example based on self-aggregation of regions carrying identical epigenetic marks or the emergence of globule structures resulting from phase separation^99–102^. Interestingly, the HCD genomic interval harbors several validated enhancers that were inactive in the embryonic heart (DE5-12). An intriguing hypothesis raised by these observations is therefore that HCDs could act to topologically sequester regulatory regions for modulation of target gene interaction in a tissue-specific manner.

From a disease perspective, our findings also expand on former analyses demonstrating that *Shox2* gene desert limb and hindbrain enhancer activities emerge within the similar-sized gene desert flanking the human *SHOX*^47,103^. Pointing to functional homology with the mouse *Shox2* regulatory region, disruption of enhancers within the gene desert downstream of *SHOX* has been associated with Leri-Weill dyschondrosteosis (LWD) and idiopathic short stature (ISS) syndromes in a significant fraction of cases^104^. Furthermore, *SHOX* haploinsufficiency is directly associated with the skeletal abnormalities observed in LWD and Turner syndrome, the latter also involving craniofacial abnormalities^105–107^. One study has also found a link between neurodevelopmental disorders and microduplications at the *SHOX* locus, suggesting that such perturbations may alter neural development or function^108^. Thus, considering the overlapping expression patterns and critical functions of human *SHOX* and mouse *Shox2*, our results provide a blueprint for the investigation of *SHOX* regulation in the hindbrain, thalamus, pharyngeal arches, and limbs^109,110^. It will be particularly interesting to determine whether “orthologous” craniofacial, neural and/or limb enhancers exist, and whether human *SHOX* enhancers share motif content or other enhancer grammar characteristics with mouse *Shox2* enhancers. Indeed, in a recent example orthologous enhancer-like sequence was identified 160kb downstream of human *SHOX* and 47kb downstream of mouse *Shox2*, respectively, and drove overlapping activities in the hindbrain^47,103^. Such enhancers presumably originate from a single ancestral *Shox* locus, preceding the duplication of *Shox* and *Shox2* paralogs and are therefore considered evolutionary ancient. Within this context, future comparative studies should include a search for deeply conserved orthologs of *SHOX* and *SHOX2* enhancers in basal chordates such as amphioxus, which express their single *Shox2* gene in the developing hindbrain^111^. The recent identification of orthologous *Islet* gene enhancers in sponges and vertebrates demonstrate the promise of such an approach^112^. Taken together, functional enhancer characterization along with refined enhancer grammar and 3D interactions at the cell type level will likely be key to resolve the regulatory complexity inherent to distributed enhancer landscapes and to understand how transcriptional dynamics and morphological complexity are rooted in gene deserts.

## MATERIALS AND METHODS

### Ethics statement and approval of animal experimentation

All animal (mouse) experiments were performed in accordance with national laws and approved by the national and local regulatory authorities. All animal work at Lawrence Berkeley National Laboratory (LBNL, CA, USA) was reviewed and approved by the LBNL Animal Welfare Committee. Knockout and transgenic mice were housed at the Animal Care Facility (the ACF) at LBNL. Mice were monitored daily for food and water intake, and animals were inspected weekly by the Chair of the Animal Welfare and Research Committee and the head of the animal facility in consultation with the veterinary staff. The LBNL ACF is accredited by the American Association for the Accreditation of Laboratory Animal Care International (AAALAC). Animal work at the University of Calgary involving the production, housing and analysis of mouse lines depicted in **Figures 6, S6 and S7**, was approved by the Life and Environmental Sciences Animal Care Committee (LESACC). All experiments with mice were performed in accordance with Canadian Council on Animal Care guidelines as approved by the University of Calgary LESACC, Protocol # AC13-0053. Animal work in Switzerland involving SV enhancer KO mice (**Figs. 5 and S5**) was approved by the regional commission on Animal Experimentation and the Cantonal Veterinary Office of the city of Bern. The following developmental stages were used in this study: embryonic day E10.5, E11.5, E12.5, E13.5 and newborn mice. Animals of both sexes were used in these analyses. Sample size selection and randomization strategies were conducted as follows:

#### Transgenic mouse assays

Sample sizes were selected empirically based on our previous experience of performing transgenic mouse assays for >3,000 total putative enhancers (VISTA Enhancer Browser: https://enhancer.lbl.gov/). Mouse embryos were excluded from further analysis if they did not encode the reporter transgene or if the developmental stage was not correct. All transgenic mice were treated with identical experimental conditions. Randomization and experimenter blinding were unnecessary and not performed.

#### Knockout mice

Sample sizes were selected empirically based on our previous studies^19,20^. All phenotypic characterization of knockout mice employed a matched littermate selection strategy. Analyzed *Shox2* gene desert and enhancer knockout embryos and mice described in this paper resulted from crossing mice heterozygous for the respective deletion to allow for the comparison of matched littermates of different genotypes. Embryonic littermates and samples from genetically modified animals were dissected and processed blind to genotype.

### Transgenic reporter analysis in mouse embryos

Transgenic reporter assays for validation of all elements except PLEs were performed at LBNL in *Mus musculus* FVB strain mice and injection of *LacZ* reporter constructs was conducted as previously described^19,113^. The related primer sequences and genomic coordinates are listed in **Tables S2 and S8**. Predicted enhancer elements were PCR-amplified from mouse genomic DNA (Clontech) and cloned into a Hsp68-*LacZ* expression vector for random integration^114^. For higher accuracy in absence of position effects, the +325 SV enhancer element was analyzed in a *β*-globin-LacZ construct for site-directed integration at the neutral *H11* locus^25,115^. PLE elements were PCR-amplified from bacterial artificial chromosomes (**Table S11**) and then cloned into the *βlacz* plasmid containing a minimal human *β*-globin promoter-*LacZ* cassette, as described^47^. Due to their large size, PLE3 (10,351 bp) and PLE5 (9,473 bp) were amplified with the proofreading polymerase in the SequalPrep^TM^ Long PCR Kit (Invitrogen). PLE transgenic mice and embryos were produced at the University of Calgary Centre for Mouse Genomics by pronuclear injection of DNA constructs into CD-1 single-cell stage embryos as described^116^. Male founder animals (or male F1 progeny produced from transgenic females) were crossed to CD-1 females to produce transgenic embryos which were stained with X-gal by standard techniques^113^.

### CRISPR/Cas9 deletion mouse lines

SgRNAs located 5’ and 3’ of the genomic sequence of interest were designed using CHOPCHOP^117^ (*Shox2* gene desert deletion) or CRISPOR^118^ (http://crispor.tefor.net/) (SV enhancer deletion). Gene desert deletion (582kb) was engineered using CRISPR/Cas9 genome editing in fertilized mouse oocytes as described, with minor modifications^114^. Briefly, a mix containing Cas9 mRNA (final concentration of 100 ng/ul) and two single guide RNAs (sgRNAs) (25 ng/ul each) was microinjected into the cytoplasm of fertilized FVB strain oocytes. The SV enhancer deletion allele was engineered using CRISPR-EZ^68^ at the Center of Transgenic Models (CTM) of the University of Basel. Genomic deletion coordinates and sgRNA sequences used for genome editing are listed in **Table S4**. Deletion alleles in Founder (F0) mice and F1 offspring were identified using PCR and the exact deletion breakpoints were verified by Sanger sequencing. Mice and embryos were PCR-genotyped using primer pairs specific for the deleted genomic interval or wild-type counterpart (**Table S5**).

### ENCODE H3K27ac ChIP-seq and mRNA-seq analysis

To establish a heatmap revealing putative enhancers and their temporal activities within the *Shox2* TAD interval, a previously generated catalog of strong enhancers identified using ChromHMM^53^ across mouse development was used^54^. Briefly, calls across 66 different tissue-stage combinations were merged and H3K27ac signals quantified as log2-transformed RPKM. Estimates of statistical significance for these signals were associated to each region for each tissue-stage combination using the corresponding H3K27ac ChIP-seq peak calls. These were downloaded from the ENCODE Data Coordination Center (DCC) (http://www.encodeproject.org/, see **Table S1**, *sheet 3* for the complete list of sample identifiers). To this purpose, short reads were aligned to the mm10 assembly of the mouse genome using bowtie (ref), with the following parameters: *-a -m 1 -n 2 -l 32 -e 3001*. Peak calling was performed using MACS v1.4^119^, with the following arguments: *--gsize=mm --bw=300 --nomodel --shiftsize=100*. Experiment-matched input DNA was used as control. Evidence from two biological replicates was combined using IDR (https://www.encodeproject.org/data-standards/terms/). The *q*-value provided in the replicated peak calls was used to annotate each putative enhancer region defined above. In case of regions overlapping more than one peak, the lowest *q*-value was used. RNA-seq raw data was downloaded from the ENCODE DCC (http://www.encodeproject.org/, see **Table S1**, *sheet 3* for the complete list of sample identifiers).

### Region Capture Hi-C (CHi-C)

Embryonic forelimbs, mandibular processes, and hearts from wildtype FVB embryos at E11.5 were micro-dissected in cold 1xPBS, pooled according to tissue type, and homogenized using a Dounce tissue grinder. Cells were resuspended in 10% FCS (in PBS) and formaldehyde (37%) diluted to a final 2% in a total volume of 1ml was added for fixation for 10 min, as previously described^60^. 1.25M Glycine was used to quench fixation and pellets were snap-frozen in liquid nitrogen and stored at −80C. Pellets were resuspended in fresh lysis buffer (10mM Tris, pH7.5, 10mM NaCl, 5mM MgCl2, 0.1 mM EGTA complemented with Protease Inhibitor) for nuclei isolation. Following 10min incubation on ice, samples were washed with 1xPBS and frozen in liquid nitrogen. 3C-libraries were prepared from thawed nuclei subjected to DpnII digestion (NEB, R0543M), re-ligated with T4 ligase (Thermo Fisher Scientific) and de-crosslinking as described previously^60^. For 3C-library quality control, 500ng of library sample along with digested and undigested control samples was assessed using agarose gel electrophoresis (1% gel). Shearing on re-ligated products was performed using a Covaris ultrasonicator (duty cycle: 10%, intensity 5, cycles per burst: 200, time: 2 cycles of 60s each). Following adaptor ligation and amplification of sheared DNA fragments, libraries were hybridized to custom-designed SureSelect beads (SureSelectXT Custom 0.5-2.9Mb library) and indexed following Agilent’s instructions. Multiplexed libraries were sequenced using 50bp paired-end sequencing (HiSeq 4000 sequencer). CHi-C probes of the SureSelect library were designed to span the *Shox2* genomic interval and adjacent TADs (mm10: chr3:65196079-68696078).

### CHi-C data processing and analysis

CHi-C processing was performed using a previously published pipeline^26^. Briefly, sequenced reads were mapped to the reference genome GRCm38/mm10 following the HiCUP pipeline^120^ (v0.8.1) set up with Bowtie2^121^ (v2.4.5). Filtering and de-duplication was conducted using HiCUP (no size selection, Nofill: 1, format: Sanger) and unique MAPQ ≥ 30 valid read pairs were obtained for FL, MD and HT datasets (N=637163, N=577862 and N=592498, respectively). Binned contact maps from valid read pairs were generated using Juicer command line tools^122^ (v1.9.9) and raw .cool files were generated with the hicConvertFormat tool (HiCExplorer v3.7.2) from native .hic out-puts generated by Juicer. For normalization and diagonal filtering the Cooler matrix balancing tool^123^ (v0.8.11) was applied with the options ‘--mad-max 5 --min-nnz 10 --min-count 0 --ignore-diags 2 -­tol 1e-05 --max-iters 200 --cis-only’. Only the targeted genomic interval enriched in the capture step (mm10: chr3:65196079-68696078) was selected for binning and balancing. Consequently, only read pairs mapping to this interval were retained, shifted by the offset of 65,196,078 bp using custom crhom.sizes files. Balanced maps were then exported at 5kb resolution with corrected coordinates (transformed back to original values). Subtraction maps were directly generated from Cooler balanced HiCmaps using hicCompareMatrices tool (HiCExplorer v3.7.2) with option ‘--operation diff’. HiCExplorer^124^ (v3.7.2) was used to determine normalized inter-domain insulation scores and domain boundaries on Hi-C and subtraction maps using default parameters ‘hicFindTADs -t 0.05 -d 0.01 -c fdr’ computing p-values for minimal window length 50000. Hi-C maps and related graphs were visualized from .cool files and bedgraph matrices, respectively, using pyGenomeTracks^125^ (*v.3.6*). GOTHiC^126^ (v.1.32.0) was used to identify reliable and significance-based Hi-C interactions from HiCUP validated read pairs (MAPQ10) with ‘res=1000, restrictionFile, cistrans=’all’, parallel=FALSE, cores=NULL’ (R pipeline-template script, v.4.2.2) and a threshold of ‘-log(q-value) > 1’.

### Virtual 4C (V4C)

To determine target interactions of a defined element locally V4C profiles were generated as described^60^ from filtered unique read pairs (hicup.bam files) which also served as input for computation of CHi-C maps (see above). Conditions for mapped read-pairs included MAPQ ≥ 30 and relative position of the two reads inside and outside the viewpoint, respectively. After quantitation of reads outside of the viewpoint (per restriction fragment), read counts were distributed into 3kb bins (with proportional distribution of read counts in case of overlap with more than one bin). Following smoothing of each binned profile via averaging^60^, peak profiles were generated using custom Java code based on htsjdk v2.12.0 (https://samtools.github.io/htsjdk/). A 10kb viewpoint containing the extended *Shox2* promoter region (chr3:66975788-66985788) was used for comparison with Hi-C maps. The viewpoint and neighboring +/-5kb regions were excluded from computation of the scaling factor. BigwigCompare tool (deepTools v3.5.1) was used to generate relative subtraction Capture-C-like profiles.

### 4C-seq from proximal forelimbs

Per replicate, 10-12 proximal forelimbs from CD-1 embryos at E12.5 were dissected in PBS, followed by 4C-seq tissue processing as described^127,128^. For tissue preparation, cells were dissociated by incubating the pooled tissue in 250μl PBS supplemented with 10% fetal fetal calf serum (FCS) and 1 mg/ml collagenase (Sigma) for 45 minutes at 37° C with shaking at 750 rpm. The solution was passed through a cell strainer (Falcon) to obtain single cells which were fixed in 9.8 ml of 2% formaldehyde in PBS/10% FCS for 10 minutes at room temperature and lysed. Libraries were prepared by overnight digestion with NlaIII (New England Biolabs (NEB)) and ligation for 4.5 hours with 100 units T4 DNA ligase (Promega, #M1794) under diluted conditions (7 ml), followed by de­crosslinking overnight at 65°C after addition of 15ul of 20mg/ml proteinase K. After phenol/chloroform extraction and ethanol precipitation the samples were digested overnight with the secondary enzyme DpnII (NEB) followed by phenol/chloroform extraction, ethanol precipitation purification and ligation for 4.5 hours in a 14 ml volume. The final ligation products were extracted and precipitated as above followed by purification using Qiagen nucleotide removal columns. For each viewpoint, libraries were prepared with 100 ng of template in each of 16 separate PCR reactions using the Roche, Expand Long Template kit with primers incorporating Illumina adapters. Viewpoint and primer details are presented in **Table S10**. PCR reactions for each viewpoint were pooled and purified with the Qiagen PCR purification kit and sequenced with the Illumina HiSeq to generate single 100bp reads. Demultiplexed reads were mapped and analyzed with the 4C-seq module of the HTSstation pipeline as described^129^. Results are shown in UCSC browser format as normalized reads per fragment after smoothing with an 11-fragment window and mapped to mm10 (**Fig. 6B**, **S6D**). Raw and processed (bedgraph) sequence files are available under GEO accession number GSE161194.

### *In situ* hybridization (ISH) and quantitative real-time PCR (qPCR)

For assessment of spatial gene expression changes in mouse embryos, whole mount *in situ* hybridization (ISH) using digoxigenin-labeled antisense riboprobes was performed as previously described^130^. At least three independent embryos were analyzed for each genotype. Embryonic tissues were imaged using a Leica MZ16 microscope coupled to a Leica DFC420 digital camera. Brightness and contrast were adjusted uniformly using Photoshop CS5. For qPCR of samples involving the gene desert deletion (GD^Δ^), isolation of RNA from micro-dissected embryonic tissues was performed using the Ambion RNAqueous Total RNA Isolation Kit (Life Technologies) according to the manufacturer’s protocol. For qPCR of samples involving the SV enhancer deletion (SV-Enh^Δ^), RNeasy Micro Kit (Qiagen) was used following the manufacturer’s protocol. RNA was reverse transcribed using SuperScript III (Life Technologies) with poly-dT priming according to manufacturer instructions. qPCR was conducted on a LightCycler 480 (Roche) using KAPA SYBR FAST qPCR Master Mix (Kapa Biosystems) for GD^Δ^ samples, and on a and on a ViiA 7 Real-Time PCR System using PowerTrack SYBR Green Master Mix (Applied Biosystems) for SV-Enh^Δ^ samples, according to manufacturer instructions. The qPCR primers used (**Table S6**) were described previously^19^. Relative gene expression levels were calculated using the 2^-aac^t method, normalized to the *Actb* housekeeping gene, and the mean of wild-type control samples was set to 1, as used previously^19^.

### Immunofluorescence (IF)

IF was performed as previously described^19^. Briefly, mouse embryos at E11.5 were isolated in cold PBS and fixed in 4% PFA for 2-3h. After incubation in a sucrose gradient and embedding in a 1:1 mixture of 30% sucrose and OCT compound, sagittal 10μm frozen tissue sections were obtained using a cryostat. Selected cryo-sections were then incubated overnight with the following primary antibodies: anti-Shox2 (1:300, Santa Cruz JK-6E, sc-81955), anti-SMA-Cy3 (1:250, Sigma, C6198), anti-Hcn4 (1:500, Thermo Fisher, MA3-903) and anti-Nkx2.5 (1:500, Thermo Fisher, PA5-81452). Goat-anti mouse, goat anti-rabbit and donkey anti-rat secondary antibodies conjugated to Alexa Fluor 488, 568, or 647 (1:1,000, Thermo Fisher Scientific) were used for detection. Hoechst 33258 (Sigma-Aldrich) was utilized to counterstain nuclei. A Zeiss AxioImager fluorescence microscope in combination with a Hamamatsu Orca-03 camera was used to acquire fluorescent images. Brightness and contrast were adjusted uniformly using Photoshop CS5.

### Skeletal preparations

Euthanized newborn mice were eviscerated, skinned and fixed in 1 % acetic acid in EtOH for 24 hours. Cartilage was stained overnight with 1 mg/mL Alcian blue 8GX (Sigma) in 20% acetic acid in EtOH. After washing in EtOH for 12 hours and treatment with 1.5 % KOH for three hours, bones were stained in 0.15 mg/mL Alizarin Red S (Sigma) in 0.5 % KOH for four hours, followed by de­staining in 20 % glycerol, 0.5 % KOH.

### X-ray micro-computed tomography (pCT) of adult mouse skeletons

Mice were euthanized at 6 weeks of age and whole-body μCT scans were generated using a Skyscan 1173 v1.6 μCT scanner (Bruker, Kontich, Belgium) at 80-85 kV and 56-62 pA with 45 gm resolution^131^ NRecon v1.7.4.2 (Bruker, Kontich, Belgium) was used to perform stack reconstructions, and 3D landmarks were placed in MeshLab^132^ (v2020.07) by one observer (CSS) blind to the genotype identity of individual animals. To quantify the length of the stylopod bones, distances were calculated between two landmarks placed at the proximal and distal ends of the humerus and femur (the proximal epiphysis [PE] and olecranon fossa lateral [OFL] for the humerus, and the greater trochanter [GT] and lateral inferior condyle [LIC] for the femur). To account for body size variability between individuals, these measurements were normalized to the inter-landmark distance between the proximal and distal ends of the third metatarsal. To assess intra-observer repeatability, CSS placed the landmarks on scans of 12 mice (six GD^Δ^/ *Shox2*^Δc^, two GD^Δ/+^, two *Shox2*^Δc/+^, and two WT) five times each, with each session separated by at least 24 hours^133^. An absolute coefficient of variation (CV) for each landmark was calculated and the average CV was 0.28% with a range of 0.14% - 0.42%.

### ATAC-seq

ATAC-seq was performed as described^134^ with minor modifications. Per replicate, pairs of wildtype mouse embryonic hearts at E11.5 were micro-dissected in cold PBS and cell nuclei were dissociated in Lysis buffer using a Dounce tissue grinder. Approx. 50’000 nuclei were then pelleted at 500 RCF for 10 min at 4°C and resuspended in 50 μL transposition reaction mix containing 25 μL Nextera 2x TD buffer and 2.5 μL TDE1 (Nextera Tn5 Transposase; Illumina) (cat. no. FC-121-1030) followed by incubation for 30 minutes at 37°C with shaking. The reaction was purified using the Qiagen MinElute PCR purification kit and amplified using defined PCR primers^135^. ATAC-seq libraries were purified using the Qiagen MinElute PCR purification kit (ID: 28004), quantified by the Qubit Fluorometer with the dsDNA HS Assay Kit (Life Technologies) and quality assessed using the Agilent Bioanalyzer high sensitivity DNA analysis assay. Libraries were pooled and sequenced using single end 50 bp reads on a HiSeq 4000 (Illumina).

### Mouse ATAC-seq and ChIP-seq data processing

Analysis of mouse ATAC-seq and reprocessing of previously published ChIP-seq datasets used in this study was performed using Adaptor trimming (trim_galore_v0.6.6) by Cutadapt (https://cutadapt.readthedocs.io/), with default parameters ‘-j 1 -e 0.1 -q 20 -O 1’ for single­end, and ‘--paired -j 1 -e 0.1 -q 20 -O 1’ for paired-end data (purging trimmed reads shorter than 20bp). For read mapping, Bowtie2^121^ (version 2.4.2) was used with parameters ‘-q --no-unal -p 8 -X2000’ (ATAC-seq) and ‘-q --no-unal -p 2’ (ChIP-seq) for both single/paired-end samples. Reads were aligned to the GRCm38/mm10 reference genome using pre-built Bowtie2 indexes from the Illumina’s iGenomes collection (http://bowtie-bio.sourceforge.net/bowtie2/). Duplicates and low-quality reads (MAPQ = 255) for both single/paired-end samples were removed using SAMtools (v1.12), with pipeline parameters ‘markdup -r’ and ‘-bh -q10’, respectively^136^. ATAC-seq peak calling was performed using MACSv2^119,137^ (v2.1.0) with p-value < 0.01 and parameters ‘-t -n -f BAM -g mm --nolambda --nomodel --shif 50 --extsize 100’ for single-end, and ‘-t -n -f BAMPE -g mm --nolambda --nomodel --shif 50 --extsize 100’ for paired-end reads. For ChIP-seq peak calling, ‘-t -c -n -f BAM -g mm’ parameters were used instead. PyGenomeTracks^125^ was used for visualization of profiles and alignment with other datasets.

### Cardiac TF motif detection

An enriched collection of position weight matrices (PWMs)^138^ was limited to motifs of TFs expressed in the developing heart at E11.5. After mapping of gene symbols to the equivalent identifiers in the Ensembl103 release using the BiomaRt v2.5.0 package (R v4.1.2)^139^, only those PWMs matching TFs expressed in E11.5 hearts were selected for analysis^140^ (ENCSR691OPQ). A mean FPKM > 2 calculated across all RNA-seq replicates was used as threshold for significant expression. This filtering resulted in a set of 576 mouse TFs. 1’376 corresponding PWMs were available for 282 of these TFs^67^ which were used for motif detection by FIMO (Find Individual Motif Occurrences)^67,141^, except for 14 that were omitted since in each case, since the match identified genome-wide was included in a larger motif within the collection (**Table S9**). FIMO v5.3.0 with a standard p-value cutoff of 10^-4^ and GC-content matched backgrounds were used for screening genomic sequence for potential TF-binding sites. Motif conservation was computed using BWTOOL v1.0^142^ based on the average of individual nucleotide PhyloP (Placental) conservation scores provided by UCSC PHAST package (http://hgdownload.cse.ucsc.edu/goldenpath/mm10/phyloP60way/).

### ChIP-seq and RNA-seq from human fetal hearts

Aspects of this study involving human heart tissue samples were reviewed and approved by the Human Subjects Committee at Lawrence Berkeley National Laboratory (Protocol Nos. 00023126 and 00022756). Fetal human heart embryonic samples at post conception week 17 (LV, RV, LA, RA) were obtained from the Human Developmental Biology Resource at Newcastle University (hdbr.org), in compliance with applicable state and federal laws and with fully informed consent. Fetal samples were transported on dry ice and stored at −80C, as reported for the LV sample generated and analyzed in a previous publication^143^. Fetal human RV, LA and RA tissue samples were processed for ChIP-seq and RNA-seq analogous to the procedure published for the fetal LV sample^143^. ChIP-seq libraries were prepared using the Illumina TruSeq library preparation kit, and pooled and sequenced (50bp single end) using a HiSeq2000 (Illumina) and processed using a previously published pipeline^19^, with minor modifications. Briefly, ChIP-seq reads were obtained following quality filtering and adaptor trimming using cutadapt_v1.1 with parameter ‘-m 25 -q 20’. Bowtie^144^ (v2.0.2.0) with parameter ‘-m 1 -v 2 -p 16’ and MACS^119^ (v1.4.2) with parameter ‘-mfold = 10,30 -nomodel -p 0.0001’ were used for read mapping (hg19) and peak calling, respectively. Duplicates were removed with SAMtools^136^. RNA-seq libraries were prepared using the TruSeq Stranded Total RNA with Ribo-Zero Human/Mouse/Rat kit (Illumina) according to manufacturer instructions. An additional purification step was used to remove remaining high molecular weight products, as published^143^. RNAseq libraries were pooled and sequenced via single end 50 bp reads on a HiSeq 4000 (Illumina) and processed as previously published, with minor modifications^19^. Briefly, RNA-seq reads were preprocessed using quality filtering and adaptor trimming with cutadapt_v1.1 (‘-m 25 -q 25’). Tophat v2.0.6 was used to align RNA-seq reads to the mouse reference genome (hg19) and the reads mapping to UCSC known genes were determined by HTSeq^145^. Normalized bigWig files were generated using bedtools (bedGraphToBigWig) and IGV browser was used for visualization of profiles.

## Data availability

Raw and processed files of the NGS datasets generated are available in the NCBI GEO database with the accession codes GSE161194 (4C-seq) and GSE232887 (super-series including C-HiC, ATAC-seq, ChIP-seq and RNA-seq data). Mouse and human genome coordinates used in this manuscript are GRCm38/mm10 and GRCh37/hg19. All relevant transgenic *in vivo* enhancer data is made available at the Vista Enhancer Browser (https://enhancer.lbl.gov). Correspondence and requests for materials should be addressed to J.C. or M.O.

## Acknowledgements

We thank L. Lopez-Delisle for sharing expertise on the use of pyGenomeTracks and Capture Hi-C analysis, G. Kelman for preliminary ATAC-seq analysis and D. Duboule for hosting and supporting 4C-Seq experiments as well as training in his laboratory. We are grateful to P. Pelczar and the members of the University of Basel Center for Transgenic Models (CTM) for generation of the mouse enhancer deletion allele and thank C. Detotto and her team at the Central Animal Facilities (CAF) of the University of Bern for excellent mouse care. We are grateful to M. Docquier and her team from the iGE3 facility for preparation and sequencing of C-HiC libraries. We thank C. Fielding at the Clara Christie Centre for Mouse Genomics for pronuclear injections conducted at the University of Calgary, to J. Theodor and J. Anderson for use of the SkyScan 1173 uCT scanner and to C. Rolian and C. Unger for help with morphometric analysis. We thank V. Rapp for cloning the +325 transgenic reporter construct. We are grateful to the members of the L.A.P. and A.V. group for technical advice and the members of the M.O. and J.C. labs for useful comments on the manuscript. This work was supported by Swiss National Science Foundation (SNSF) grant PCEFP3_186993 (to M.O.), a Discovery Grant (RGPIN/355731-2013) from the Natural Sciences and Engineering Research Council of Canada (to J.C.) and National Institutes of Health grants R01HG003988, U54HG006997, R24HL123879 and UM1HL098166 (to A.V. and L.A.P.). MO was also supported by grants of the Swiss Heart Foundation (FF20110) and Novartis Foundation for Medical-Biological Research (#21C183). J.L-R. is supported by the MICINN grants PID2020-113497GB-I00 and CEX2020-001088-M (Unidad de Excelencia María de Maeztu institutional grant). G.A. is supported by Swiss National Science Foundation Grant PP00P3_176802. F.D. is supported by a SNSF postdoc.mobility fellowship (P400PB_194334). Research at the E.O. Lawrence Berkeley National Laboratory was performed under Department of Energy Contract DE-AC02-05CH11231, University of California.

## Author contributions

M.O. and J.C. conceived the study. S.A.-O., B.J.M, and M.Z. performed critical experimental (S.A.-O., B.J.M.) and computational (M.Z.) analyses for the study. S.A.-O., B.J.M., J.C. and M.O. designed and performed transgenic reporter and gene expression analyses. R.R. conducted experimental C­HiC. V.T. and J.L.-R. executed the *in-situ* hybridization analysis. M.Z. performed C-HiC and ATAC-seq/ChIP-seq data processing and analysis from all mouse datasets. I.B. set up the enhancer profiling framework based on ENCODE data and ChromHMM. C.H.S. and B.J.M. conducted ChIP-seq and RNA-seq from human heart tissues. Y. F.-Y. performed ChIP-seq and RNA-seq processing and analysis of human heart datasets. S.A.-O., E.R-C., A.L., G.A. and J.C. performed 4C-seq experiments and analysis. V.R. and J.G. conducted SV enhancer-deletion experiments. F.D., A.L., R.H., J.A.A. performed additional experimental work related to transgenic reporter validation. T.A.F and C.S.S. did skeletal phenotyping. C.S.N, I.P.-F. and S.T. performed pro-nuclear injections. G.A., D.E.D., A.V. and L.A.P. provided project funding and support. J.C. and M.O. provided project funding and wrote the manuscript with input from the other authors.

## REFERENCES

1. Craig Venter, J., et al. The Sequence of the Human Genome. Science 291, 1304–1351 (2001).

2. Ovcharenko, I. et al. Evolution and functional classification of vertebrate gene deserts. Genome Res. 15, 137–145 (2005).

3. Nobrega, M. A., Ovcharenko, I., Afzal, V. & Rubin, E. M. Scanning human gene deserts for long-range enhancers. Science 302, 413 (2003).

4. Catarino, R. R. & Stark, A. Assessing sufficiency and necessity of enhancer activities for gene expression and the mechanisms of transcription activation. Genes Dev. 32, 202–223 (2018).

5. Nobrega, M. A., Zhu, Y., Plajzer-Frick, I., Afzal, V. & Rubin, E. M. Megabase deletions of gene deserts result in viable mice. Nature 431, 984–988 (2004).

6. Montavon, T. et al. A regulatory archipelago controls Hox genes transcription in digits. Cell 147, 1132–1145 (2011).

7. Andrey, G. et al. A switch between topological domains underlies HoxD genes collinearity in mouse limbs. Science 340, 1234167 (2013).

8. Rodriguez-Carballo, E. et al. The HoxD cluster is a dynamic and resilient TAD boundary controlling the segregation of antagonistic regulatory landscapes. Genes Dev. 31, 2264–2281 (2017).

9. Darbellay, F. & Duboule, D. Topological Domains, Metagenes, and the Emergence of Pleiotropic Regulations at Hox Loci. Curr. Top. Dev. Biol. 116, 299–314 (2016).

10. Franke, M. et al. Formation of new chromatin domains determines pathogenicity of genomic duplications. Nature 538, 265–269 (2016).

11. Symmons, O. et al. The Shh Topological Domain Facilitates the Action of Remote Enhancers by Reducing the Effects of Genomic Distances. Dev. Cell 39, 529–543 (2016).

12. Marinic, M., Aktas, T., Ruf, S. & Spitz, F. An integrated holo-enhancer unit defines tissue and gene specificity of the Fgf8 regulatory landscape. Dev. Cell 24, 530–542 (2013).

13. Schoenfelder, S. & Fraser, P. Long-range enhancer-promoter contacts in gene expression control. Nat. Rev. Genet. 20, 437–455 (2019).

14. Furlong, E. E. M. & Levine, M. Developmental enhancers and chromosome topology. Science 361, 1341–1345 (2018).

15. Sanborn, A. L. et al. Chromatin extrusion explains key features of loop and domain formation in wild-type and engineered genomes. Proc. Natl. Acad. Sci. U. S. A. 112, E6456–65 (2015).

16. Fudenberg, G. et al. Formation of Chromosomal Domains by Loop Extrusion. Cell Rep. 15, 2038–2049 (2016).

17. Lupianez, D. G. et al. Disruptions of topological chromatin domains cause pathogenic rewiring of gene-enhancer interactions. Cell 161, 1012–1025 (2015).

18. Spielmann, M., Lupiáñez, D. G. & Mundlos, S. Structural variation in the 3D genome. Nat. Rev. Genet. 19, 453–467 (2018).

19. Osterwalder, M. et al. Enhancer redundancy provides phenotypic robustness in mammalian development. Nature 554, 239–243 (2018).

20. Dickel, D. E. et al. Ultraconserved Enhancers Are Required for Normal Development. Cell 172, 491–499.e15 (2018).

21. Hornblad, A., Bastide, S., Langenfeld, K., Langa, F. & Spitz, F. Dissection of the Fgf8 regulatory landscape by in vivo CRISPR-editing reveals extensive intra- and inter-enhancer redundancy. Nat. Commun. 12, 439 (2021).

22. Will, A. J. et al. Composition and dosage of a multipartite enhancer cluster control developmental expression of Ihh (Indian hedgehog). Nat. Genet. 49, 1539–1545 (2017).

23. van Mierlo, G., Pushkarev, O., Kribelbauer, J. F. & Deplancke, B. Chromatin modules and their implication in genomic organization and gene regulation. Trends Genet. (2022) doi:10.1016/j.tig.2022.11.003.

24. Malkmus, J. et al. Spatial regulation by multiple Gremlin1 enhancers provides digit development with cis-regulatory robustness and evolutionary plasticity. Nat. Commun. 12, 5557 (2021).

25. Kvon, E. Z. et al. Comprehensive In Vivo Interrogation Reveals Phenotypic Impact of Human Enhancer Variants. Cell 180, 1262–1271.e15 (2020).

26. Rouco, R. et al. Cell-specific alterations in Pitx1 regulatory landscape activation caused by the loss of a single enhancer. Nat. Commun. 12, 7235 (2021).

27. Cobb, J., Dierich, A., Huss-Garcia, Y. & Duboule, D. A mouse model for human short-stature syndromes identifies Shox2 as an upstream regulator of Runx2 during long-bone development. Proc. Natl. Acad. Sci. U. S. A. 103, 4511–4515 (2006).

28. Gu, S., Wei, N., Yu, L., Fei, J. & Chen, Y. Shox2-deficiency leads to dysplasia and ankylosis of the temporomandibular joint in mice. Mech. Dev. 125, 729–742 (2008).

29. Yu, L. et al. Shox2-deficient mice exhibit a rare type of incomplete clefting of the secondary palate. Development 132, 4397–4406 (2005).

30. Rosin, J. M., Kurrasch, D. M. & Cobb, J. Shox2 is required for the proper development of the facial motor nucleus and the establishment of the facial nerves. BMC Neurosci. 16, 39 (2015).

31. Rosin, J. M. et al. Mice lacking the transcription factor SHOX2 display impaired cerebellar development and deficits in motor coordination. Dev. Biol. 399, 54–67 (2015).

32. Scott, A. et al. Transcription factor short stature homeobox 2 is required for proper development of tropomyosin-related kinase B-expressing mechanosensory neurons. J. Neurosci. 31, 6741–6749 (2011).

33. Xu, J. et al. Shox2 regulates osteogenic differentiation and pattern formation during hard palate development in mice. J. Biol. Chem. 294, 18294–18305 (2019).

34. Blaschke, R. J. et al. Targeted mutation reveals essential functions of the homeodomain transcription factor Shox2 in sinoatrial and pacemaking development. Circulation 115, 1830–1838 (2007).

35. Espinoza-Lewis, R. A. et al. Shox2 is essential for the differentiation of cardiac pacemaker cells by repressing Nkx2-5. Dev. Biol. 327, 376–385 (2009).

36. van Eif, V. W. W. et al. Transcriptome analysis of mouse and human sinoatrial node cells reveals a conserved genetic program. Development 146, (2019).

37. Ye, W. et al. A common Shox2-Nkx2-5 antagonistic mechanism primes the pacemaker cell fate in the pulmonary vein myocardium and sinoatrial node. Development 142, 2521–2532 (2015).

38. Hoffmann, S. et al. Coding and non-coding variants in the SHOX2 gene in patients with early-onset atrial fibrillation. Basic Res. Cardiol. 111, 36 (2016).

39. Hoffmann, S. et al. Functional Characterization of Rare Variants in the SHOX2 Gene Identified in Sinus Node Dysfunction and Atrial Fibrillation. Front. Genet. 10, 648 (2019).

40. Li, N. et al. A SHOX2 loss-of-function mutation underlying familial atrial fibrillation. Int. J. Med. Sci. 15, 1564–1572 (2018).

41. Mori, A. D. et al. Tbx5-dependent rheostatic control of cardiac gene expression and morphogenesis. Dev. Biol. 297, 566–586 (2006).

42. Vedantham, V., Galang, G., Evangelista, M., Deo, R. C. & Srivastava, D. RNA sequencing of mouse sinoatrial node reveals an upstream regulatory role for Islet-1 in cardiac pacemaker cells. Circ. Res. 116, 797–803 (2015).

43. Puskaric, S. et al. Shox2 mediates Tbx5 activity by regulating Bmp4 in the pacemaker region of the developing heart. Hum. Mol. Genet. 19, 4625–4633 (2010).

44. Galang, G. et al. ATAC-Seq Reveals an Isl1 Enhancer That Regulates Sinoatrial Node Development and Function. Circ. Res. 127, 1502–1518 (2020).

45. Hoffmann, S. et al. Islet1 is a direct transcriptional target of the homeodomain transcription factor Shox2 and rescues the Shox2-mediated bradycardia. Basic Res. Cardiol. 108, 339 (2013).

46. Marchini, A., Ogata, T. & Rappold, G. A. A Track Record on SHOX: From Basic Research to Complex Models and Therapy. Endocr. Rev. 37, 417–448 (2016).

47. Rosin, J. M., Abassah-Oppong, S. & Cobb, J. Comparative transgenic analysis of enhancers from the human SHOX and mouse Shox2 genomic regions. Hum. Mol. Genet. 22, 3063–3076 (2013).

48. Liu, H., Jiao, Z., Espinoza-Lewis, R. A., Chen, C. & Chen, Y. FUNCTIONAL REDUNDANCY BETWEEN HUMAN SHOX AND MOUSE SHOX2 IN THE REGULATION OF SINUS NODE FORMATION. J. Am. Coll. Cardiol. 57, E53 (2011).

49. Ye, W. et al. A unique stylopod patterning mechanism by Shox2-controlled osteogenesis. Development 143, 2548–2560 (2016).

50. Cazalla, D., Newton, K. & Cáceres, J. F. A novel SR-related protein is required for the second step of Pre-mRNA splicing. Mol. Cell. Biol. 25, 2969–2980 (2005).

51. Scala, M. et al. RSRC1 loss-of-function variants cause mild to moderate autosomal recessive intellectual disability. Brain 143, e31 (2020).

52. Visel, A., Minovitsky, S., Dubchak, I. & Pennacchio, L. A. VISTA Enhancer Browser--a database of tissue-specific human enhancers. Nucleic Acids Res. 35, D88–92 (2007).

53. Ernst, J. & Kellis, M. ChromHMM: automating chromatin-state discovery and characterization. Nat. Methods 9, 215–216 (2012).

54. Gorkin, D. U. et al. An atlas of dynamic chromatin landscapes in mouse fetal development. Nature 583, 744–751 (2020).

55. Rodriguez-Carballo, E., Lopez-Delisle, L., Yakushiji-Kaminatsui, N., Ullate-Agote, A. & Duboule, D. Impact of genome architecture on the functional activation and repression of Hox regulatory landscapes. BMC Biol. 17, 55 (2019).

56. ENCODE Project Consortium et al. Expanded encyclopaedias of DNA elements in the human and mouse genomes. Nature 583, 699–710 (2020).

57. Mendez-Maldonado, K., Vega-López, G. A., Aybar, M. J. & Velasco, I. Neurogenesis From Neural Crest Cells: Molecular Mechanisms in the Formation of Cranial Nerves and Ganglia. Front Cell Dev Biol 8, 635 (2020).

58. Andrey, G. et al. Characterization of hundreds of regulatory landscapes in developing limbs reveals two regimes of chromatin folding. Genome Res. 27, 223–233 (2017).

59. Kragesteen, B. K. et al. Dynamic 3D chromatin architecture contributes to enhancer specificity and limb morphogenesis. Nat. Genet. 50, 1463–1473 (2018).

60. Paliou, C. et al. Preformed chromatin topology assists transcriptional robustness of Shh during limb development. Proc. Natl. Acad. Sci. U. S. A. 116, 12390–12399 (2019).

61. Bonev, B. et al. Multiscale 3D Genome Rewiring during Mouse Neural Development. Cell 171, 557–572.e24 (2017).

62. van Eif, V. W. W., Devalla, H. D., Boink, G. J. J. & Christoffels, V. M. Transcriptional regulation of the cardiac conduction system. Nat. Rev. Cardiol. 15, 617–630 (2018).

63. van Eif, V. W. W. et al. Genome-Wide Analysis Identifies an Essential Human TBX3 Pacemaker Enhancer. Circ. Res. 127, 1522–1535 (2020).

64. Fernandez-Perez, A. et al. Hand2 Selectively Reorganizes Chromatin Accessibility to Induce Pacemaker-like Transcriptional Reprogramming. Cell Rep. 27, 2354–2369.e7 (2019).

65. Akerberg, B. N. et al. A reference map of murine cardiac transcription factor chromatin occupancy identifies dynamic and conserved enhancers. Nat. Commun. 10, 4907 (2019).

66. He, A. et al. Dynamic GATA4 enhancers shape the chromatin landscape central to heart development and disease. Nat. Commun. 5, 4907 (2014).

67. Monti, R. et al. Limb-Enhancer Genie: An accessible resource of accurate enhancer predictions in the developing limb. PLoS Comput. Biol. 13, e1005720 (2017).

68. Chen, S., Lee, B., Lee, A. Y.-F., Modzelewski, A. J. & He, L. Highly Efficient Mouse Genome Editing by CRISPR Ribonucleoprotein Electroporation of Zygotes. J. Biol. Chem. 291, 14457–14467 (2016).

69. Yu, L. et al. Shox2 is required for chondrocyte proliferation and maturation in proximal limb skeleton. Dev. Biol. 306, 549–559 (2007).

70. Bobick, B. E. & Cobb, J. Shox2 regulates progression through chondrogenesis in the mouse proximal limb. J. Cell Sci. 125, 6071–6083 (2012).

71. Neufeld, S. J., Wang, F. & Cobb, J. Genetic interactions between Shox2 and Hox genes during the regional growth and development of the mouse limb. Genetics 198, 1117–1126 (2014).

72. Logan, M. et al. Expression of Cre Recombinase in the developing mouse limb bud driven by a Prxl enhancer. Genesis 33, 77–80 (2002).

73. Touceda-Suárez, M. et al. Ancient Genomic Regulatory Blocks Are a Source for Regulatory Gene Deserts in Vertebrates after Whole-Genome Duplications. Mol. Biol. Evol. 37, 2857–2864 (2020).

74. Long, H. K. et al. Loss of Extreme Long-Range Enhancers in Human Neural Crest Drives a Craniofacial Disorder. Cell Stem Cell 27, 765–783.e14 (2020).

75. de Laat, W. & Duboule, D. Topology of mammalian developmental enhancers and their regulatory landscapes. Nature 502, 499–506 (2013).

76. Lonfat, N., Montavon, T., Darbellay, F., Gitto, S. & Duboule, D. Convergent evolution of complex regulatory landscapes and pleiotropy at Hox loci. Science 346, 1004–1006 (2014).

77. Pang, B., van Weerd, J. H., Hamoen, F. L. & Snyder, M. P. Identification of non-coding silencer elements and their regulation of gene expression. Nat. Rev. Mol. Cell Biol. (2022) doi:10.1038/s41580-022-00549-9.

78. Pachano, T., Haro, E. & Rada-Iglesias, A. Enhancer-gene specificity in development and disease. Development 149, (2022).

79. Batut, P. J. et al. Genome organization controls transcriptional dynamics during development. Science 375, 566–570 (2022).

80. Statello, L., Guo, C.-J., Chen, L.-L. & Huarte, M. Gene regulation by long non-coding RNAs and its biological functions. Nat. Rev. Mol. Cell Biol. 22, 96–118 (2021).

81. Dickel, D. E. et al. Genome-wide compendium and functional assessment of in vivo heart enhancers. Nat. Commun. 7, 12923 (2016).

82. Claringbould, A. & Zaugg, J. B. Enhancers in disease: molecular basis and emerging treatment strategies. Trends Mol. Med. 27, 1060–1073 (2021).

83. van der Lee, R., Correard, S. & Wasserman, W. W. Deregulated Regulators: Disease­Causing cis Variants in Transcription Factor Genes. Trends Genet. 36, 523–539 (2020).

84. Corradin, O. & Scacheri, P. C. Enhancer variants: evaluating functions in common disease. Genome Med. 6, 85 (2014).

85. Sun, C., Zhang, T., Liu, C., Gu, S. & Chen, Y. Generation of Shox2-Cre allele for tissue specific manipulation of genes in the developing heart, palate, and limb. Genesis 51, 515–522 (2013).

86. Rada-Iglesias, A. et al. A unique chromatin signature uncovers early developmental enhancers in humans. Nature 470, 279–283 (2011).

87. Nord, A. S. et al. Rapid and pervasive changes in genome-wide enhancer usage during mammalian development. Cell 155, 1521–1531 (2013).

88. Gasperini, M., Tome, J. M. & Shendure, J. Towards a comprehensive catalogue of validated and target-linked human enhancers. Nat. Rev. Genet. 21, 292–310 (2020).

89. Mannion, B. J. et al. Uncovering Hidden Enhancers Through Unbiased In Vivo Testing. bioRxiv 2022.05.29.493901 (2022) doi:10.1101/2022.05.29.493901.

90. Kvon, E. Z., Waymack, R., Gad, M. & Wunderlich, Z. Enhancer redundancy in development and disease. Nat. Rev. Genet. 22, 324–336 (2021).

91. van Ouwerkerk, A. F. et al. Patient-Specific TBX5-G125R Variant Induces Profound Transcriptional Deregulation and Atrial Dysfunction. Circulation 145, 606–619 (2022).

92. Zhang, M. et al. Long-range Pitx2c enhancer-promoter interactions prevent predisposition to atrial fibrillation. Proc. Natl. Acad. Sci. U. S. A. 116, 22692–22698 (2019).

93. Frankel, N. et al. Phenotypic robustness conferred by apparently redundant transcriptional enhancers. Nature 466, 490–493 (2010).

94. Perry, M. W., Boettiger, A. N., Bothma, J. P. & Levine, M. Shadow enhancers foster robustness of Drosophila gastrulation. Curr. Biol. 20, 1562–1567 (2010).

95. Cannavo, E. et al. Shadow Enhancers Are Pervasive Features of Developmental Regulatory Networks. Curr. Biol. 26, 38–51 (2016).

96. Bolt, C. C. & Duboule, D. The regulatory landscapes of developmental genes. Development 147, (2020).

97. Cova, G. et al. Combinatorial effects on gene expression at the Lbx1/Fgf8 locus resolve split-hand/foot malformation type 3. Nat. Commun. 14, 1475 (2023).

98. Berlivet, S. et al. Clustering of tissue-specific sub-TADs accompanies the regulation of HoxA genes in developing limbs. PLoS Genet. 9, e1004018 (2013).

99. Rao, S. S. P. et al. A 3D map of the human genome at kilobase resolution reveals principles of chromatin looping. Cell 159, 1665–1680 (2014).

100. Rowley, M. J. & Corces, V. G. Organizational principles of 3D genome architecture. Nat. Rev. Genet. 19, 789–800 (2018).

101. Conte, M. et al. Polymer physics indicates chromatin folding variability across single-cells results from state degeneracy in phase separation. Nat. Commun. 11, 3289 (2020).

102. Chang, L.-H., Ghosh, S. & Noordermeer, D. TADs and Their Borders: Free Movement or Building a Wall? J. Mol. Biol. 432, 643–652 (2020).

103. Skuplik, I. et al. Identification of a limb enhancer that is removed by pathogenic deletions downstream of the SHOX gene. Sci. Rep. 8, 14292 (2018).

104. Chen, J., et al. Enhancer deletions of the SHOX gene as a frequent cause of short stature: the essential role of a 250 kb downstream regulatory domain. J. Med. Genet. 46, 834–839 (2009).

105. Shears, D. J., et al. Mutation and deletion of the pseudoautosomal gene SHOX cause Leri-Weill dyschondrosteosis. Nat. Genet. 19, 70–73 (1998).

106. Rappold, G. A., Shanske, A. & Saenger, P. All shook up by SHOX deficiency. The Journal of pediatrics vol. 147 422–424 (2005).

107. Rao, E., et al. Pseudoautosomal deletions encompassing a novel homeobox gene cause growth failure in idiopathic short stature and Turner syndrome. Nat. Genet. 16, 54–63 (1997).

108. Tropeano, M., et al. Microduplications at the pseudoautosomal SHOX locus in autism spectrum disorders and related neurodevelopmental conditions. J. Med. Genet. 53, 536–547 (2016).

109. Clement-Jones, M., et al. The short stature homeobox gene SHOX is involved in skeletal abnormalities in Turner syndrome. Hum. Mol. Genet. 9, 695–702 (2000).

110. Durand, C., et al. Alternative splicing and nonsense-mediated RNA decay contribute to the regulation of SHOX expression. PLoS One 6, e18115 (2011).

111. Jackman, W. R. & Kimmel, C. B. Coincident iterated gene expression in the amphioxus neural tube. Evol. Dev. 4, 366–374 (2002).

112. Wong, E. S., et al. Deep conservation of the enhancer regulatory code in animals. Science 370, (2020).

113. Kothary, R., et al. Inducible expression of an hsp68-lacZ hybrid gene in transgenic mice. Development 105, 707–714 (1989).

114. Osterwalder, M., et al. Characterization of Mammalian In Vivo Enhancers Using Mouse Transgenesis and CRISPR Genome Editing. Methods Mol. Biol. 2403, 147–186 (2022).

115. Darbellay, F. et al. Chondrogenic Enhancer Landscape of Limb and Axial Skeleton Development. bioRxiv 2023.05.10.539849 (2023) doi:10.1101/2023.05.10.539849.

116. Andras Nagy, M. G., Vintersten, K., and Behringer, R. Manipulating the Mouse Embryo: A Laboratory Manual, 3rd Edn. Cold Spring Harbor, NY: Cold Spring Harbor Laboratory Press (2003).

117. Labun, K., et al. CHOPCHOP v3: expanding the CRISPR web toolbox beyond genome editing. Nucleic Acids Res. 47, W171–W174 (2019).

118. Concordet, J.-P. & Haeussler, M. CRISPOR: intuitive guide selection for CRISPR/Cas9 genome editing experiments and screens. Nucleic Acids Res. 46, W242–W245 (2018).

119. Zhang, Y., et al. Model-based analysis of ChIP-Seq (MACS). Genome Biol. 9, R137 (2008).

120. Wingett, S., et al. HiCUP: pipeline for mapping and processing Hi-C data. F1000Res. 4, 1310 (2015).

121. Langmead, B. & Salzberg, S. L. Fast gapped-read alignment with Bowtie 2. Nat. Methods 9, 357–359 (2012).

122. Durand, N. C., et al. Juicer Provides a One-Click System for Analyzing Loop-Resolution Hi-C Experiments. Cell Syst 3, 95–98 (2016).

123. Abdennur, N. & Mirny, L. A. Cooler: scalable storage for Hi-C data and other genomically labeled arrays. Bioinformatics 36, 311–316 (2020).

124. Wolff, J., Backofen, R. & Grüning, B. Loop detection using Hi-C data with HiCExplorer. Gigascience 11, (2022).

125. Lopez-Delisle, L., et al. pyGenomeTracks: reproducible plots for multivariate genomic datasets. Bioinformatics 37, 422–423 (2021).

126. Mifsud, B., et al. GOTHiC, a probabilistic model to resolve complex biases and to identify real interactions in Hi-C data. PLoS One 12, e0174744 (2017).

127. Noordermeer, D., et al. The dynamic architecture of Hox gene clusters. Science 334, 222–225 (2011).

128. Noordermeer, D., et al. Temporal dynamics and developmental memory of 3D chromatin architecture at Hox gene loci. Elife 3, e02557 (2014).

129. David, F. P. A., et al. HTSstation: a web application and open-access libraries for high-throughput sequencing data analysis. PLoS One 9, e85879 (2014).

130. Tissieres, V., et al. Gene Regulatory and Expression Differences between Mouse and Pig Limb Buds Provide Insights into the Evolutionary Emergence of Artiodactyl Traits. Cell Rep. 31, 107490 (2020).

131. Unger, C. M., Devine, J., Hallgrímsson, B. & Rolian, C. Selection for increased tibia length in mice alters skull shape through parallel changes in developmental mechanisms. Elife 10, (2021).

132. Cignoni, P., et al. MeshLab: an Open-Source Mesh Processing Tool. Sixth Eurographics Italian Chapter Conference 129–136 (2008).

133. Cosman, M. N., Sparrow, L. M. & Rolian, C. Changes in shape and cross-sectional geometry in the tibia of mice selectively bred for increases in relative bone length. J. Anat. 228, 940–951 (2016).

134. Buenrostro, J. D., Wu, B., Chang, H. Y. & Greenleaf, W. J. ATAC-seq: A Method for Assaying Chromatin Accessibility Genome-Wide. Curr. Protoc. Mol. Biol. 109, 21.29.1-21.29.9 (2015).

135. Buenrostro, J. D., Giresi, P. G., Zaba, L. C., Chang, H. Y. & Greenleaf, W. J. Transposition of native chromatin for fast and sensitive epigenomic profiling of open chromatin, DNA-binding proteins and nucleosome position. Nat. Methods 10, 1213–1218 (2013).

136. Li, H., et al. The Sequence Alignment/Map format and SAMtools. Bioinformatics 25, 2078–2079 (2009).

137. Feng, J., Liu, T., Qin, B., Zhang, Y. & Liu, X. S. Identifying ChIP-seq enrichment using MACS. Nat. Protoc. 7, 1728–1740 (2012).

138. Diaferia, G. R., et al. Dissection of transcriptional and cis-regulatory control of differentiation in human pancreatic cancer. EMBO J. 35, 595–617 (2016).

139. Erwin, G. D., et al. Integrating diverse datasets improves developmental enhancer prediction. PLoS Comput. Biol. 10, e1003677 (2014).

140. Rahmanian, S., et al. Dynamics of microRNA expression during mouse prenatal development. Genome Res. 29, 1900–1909 (2019).

141. Grant, C. E., Bailey, T. L. & Noble, W. S. FIMO: scanning for occurrences of a given motif. Bioinformatics 27, 1017–1018 (2011).

142. Pohl, A. & Beato, M. bwtool: a tool for bigWig files. Bioinformatics 30, 1618–1619 (2014).

143. Spurrell, C. H., et al. Genome-wide fetalization of enhancer architecture in heart disease. Cell Rep. 40, 111400 (2022).

144. Langmead, B., Trapnell, C., Pop, M. & Salzberg, S. L. Ultrafast and memory-efficient alignment of short DNA sequences to the human genome. Genome Biol. 10, R25 (2009).

145. Anders, S., Pyl, P. T. & Huber, W. HTSeq--a Python framework to work with high-throughput sequencing data. Bioinformatics 31, 166–169 (2015).

