## Supplemental Figures and Tables for "A gene desert required for regulatory control of pleiotropic *Shox2* expression and embryonic survival"

### SUPPLEMENTARY FIGURES AND FIGURE LEGENDS

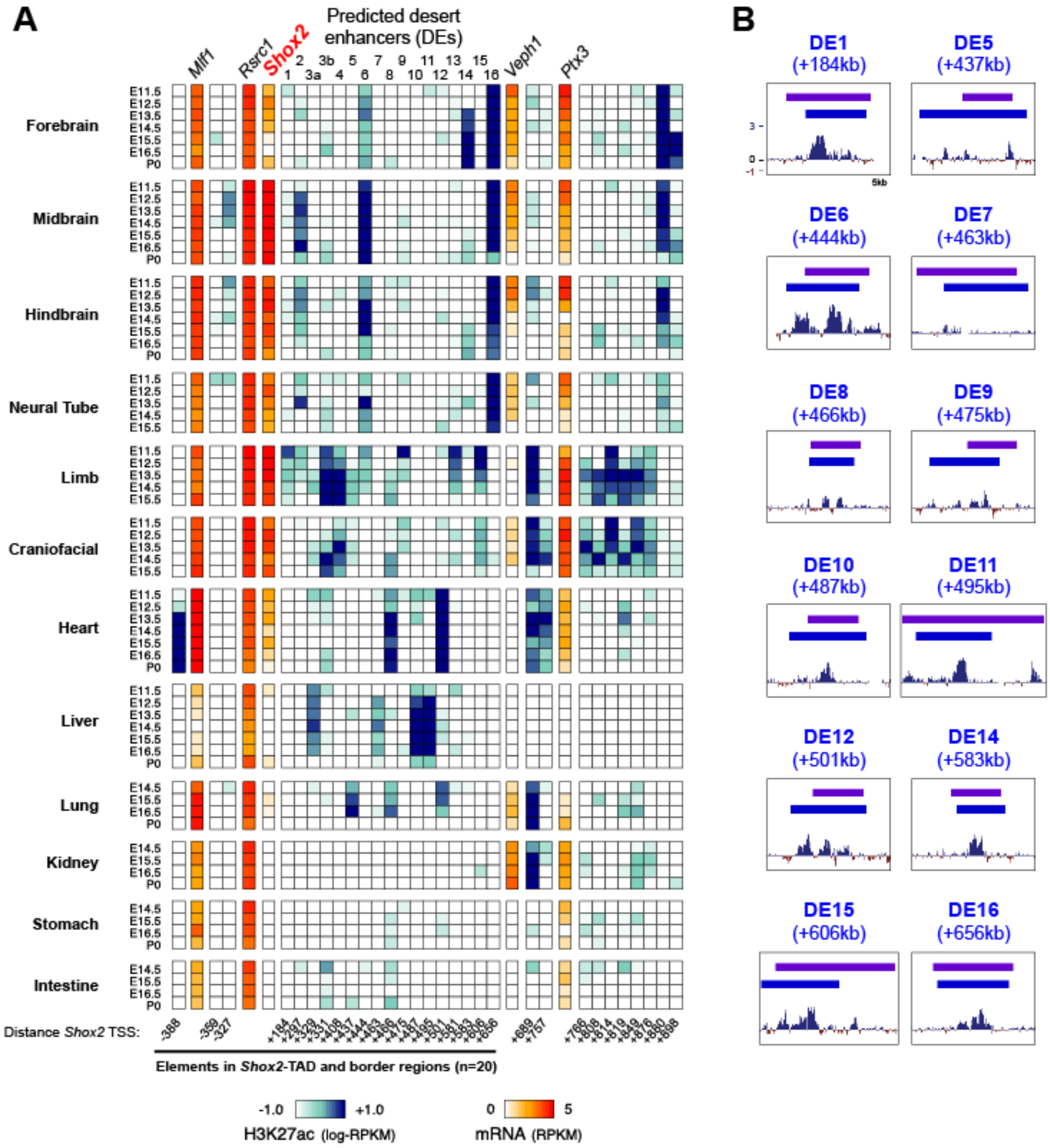


**Figure S1. Spatio-temporal H3K27ac signatures and conservation of gene desert enhancers (DEs) at the *Shox2* locus.** (**A**) Heat map for prediction of tissue-specific enhancers within the extended *Shox2* TAD[1] across different time-points and tissues, based on H3K27ac marks[2] (ENCODE) and ChromHMM filtering (see ***Methods***). Blue shading indicates levels of H3K27ac ChIP-seq enrichment and red shades illustrate transcript levels (ENCODE RNA-seq datasets) of genes located in the region (**Table S1**). Predicted gene desert enhancer (DE) IDs are listed on top with distance (in kb) to *Shox2* TSS in the bottom (-, upstream; +, downstream). (**B**) Base-wise conservation track by PhyloP (placental mammals) for each DE with validated tissue-specific enhancer activity in mouse embryos at E11.5 (**Fig. 1C**).

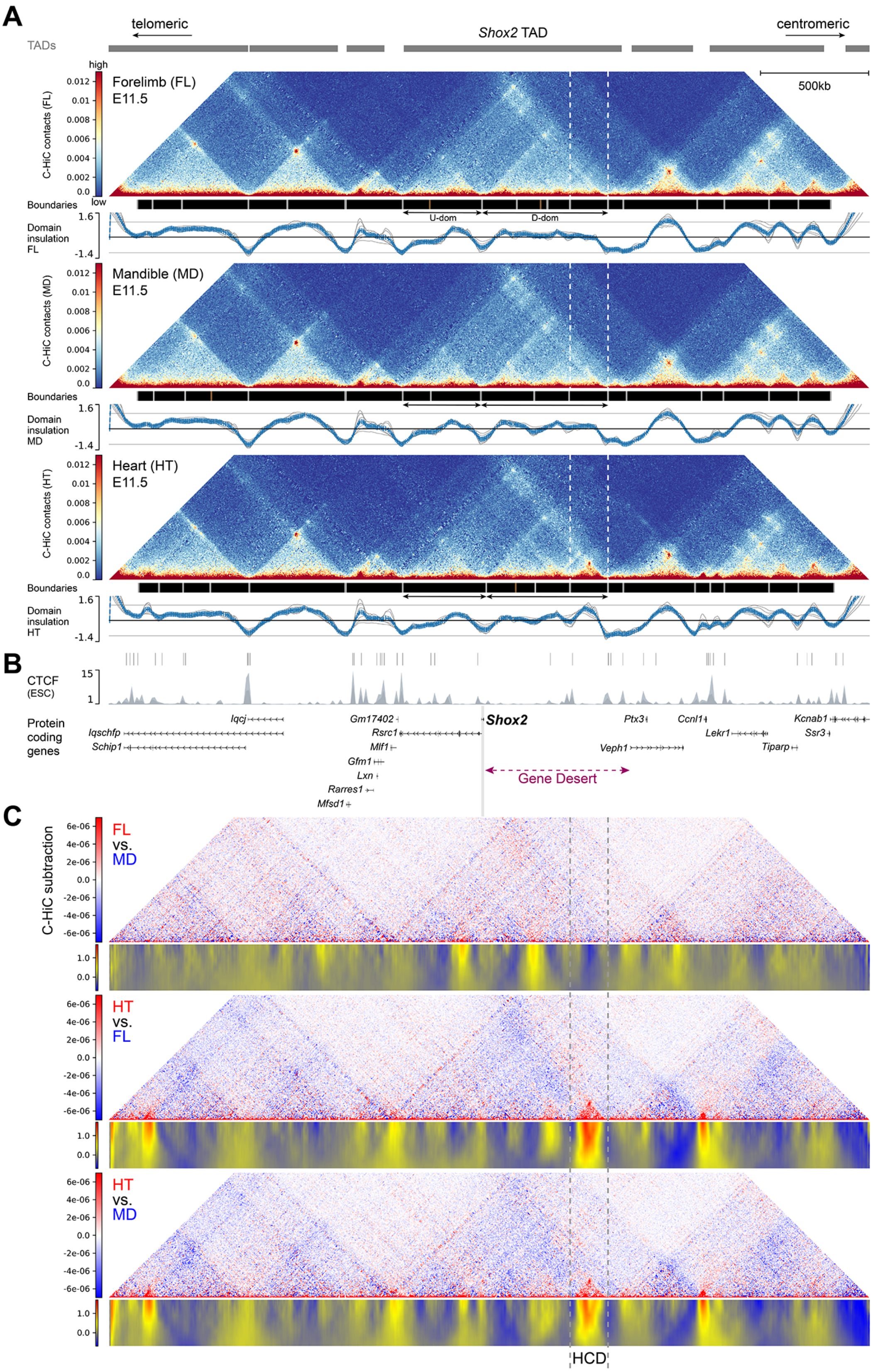


56 **Figure S2. 3D chromatin interactions across the extended *Shox2* genomic locus.** (**A**) Chromatin
57 conformation across the 3.5Mb C-HiC probe interval (mm10, chr3:65196078-68696078) centered on the
58 *Shox2* TAD from forelimb (FL), mandibular process (MD) and heart (HT) tissues of mouse wildtype embryos
59 at E11.5. Top: C-HiC matrices revealing 3D chromatin contacts. Middle: Gray boxes (with white frames) on
60 black background represent domain boundaries according to significant TAD separation score ($p<0.01$).

Brown boxes represent weaker boundaries ($p<0.05$). Bottom: Plot depicting normalized inter-domain insulation scores (lowest values correspond to strongest insulation). TAD intervals from mouse embryonic stem cells (mESCs) are represented as black bars on top[3]. *Shox2* U-dom (upstream domain) and D-Dom (downstream domain) are indicated by double arrows (**B**) Subtraction analysis identifies the HCD as a unique heart-specific >100kb chromatin domain structure in the extended genomic interval analyzed by C-HiC (see also **Fig. 2**). For each tissue, C-HiC subtraction profiles from pairwise tissue comparisons are shown on top. Red and blue shades represent tissue-enriched contacts in respective comparisons. (**B**) Top: CTCF profiles and peak calls in mESCs[3], and protein coding genes. The extension of the *Shox2* gene desert is indicated (purple double arrow). (**C**) Top: Subtraction of C-HiC profiles illustrates tissue-specific contacts in pairwise tissue comparisons as indicated (red/blue). Bottom: Matrices depicting subtraction of normalized inter-domain insulation scores based on pairwise tissue comparisons above. Strong interaction (red) is observed across the HCD interval in HT vs. FL and HT vs. MD, but not FL vs. MD comparisons.

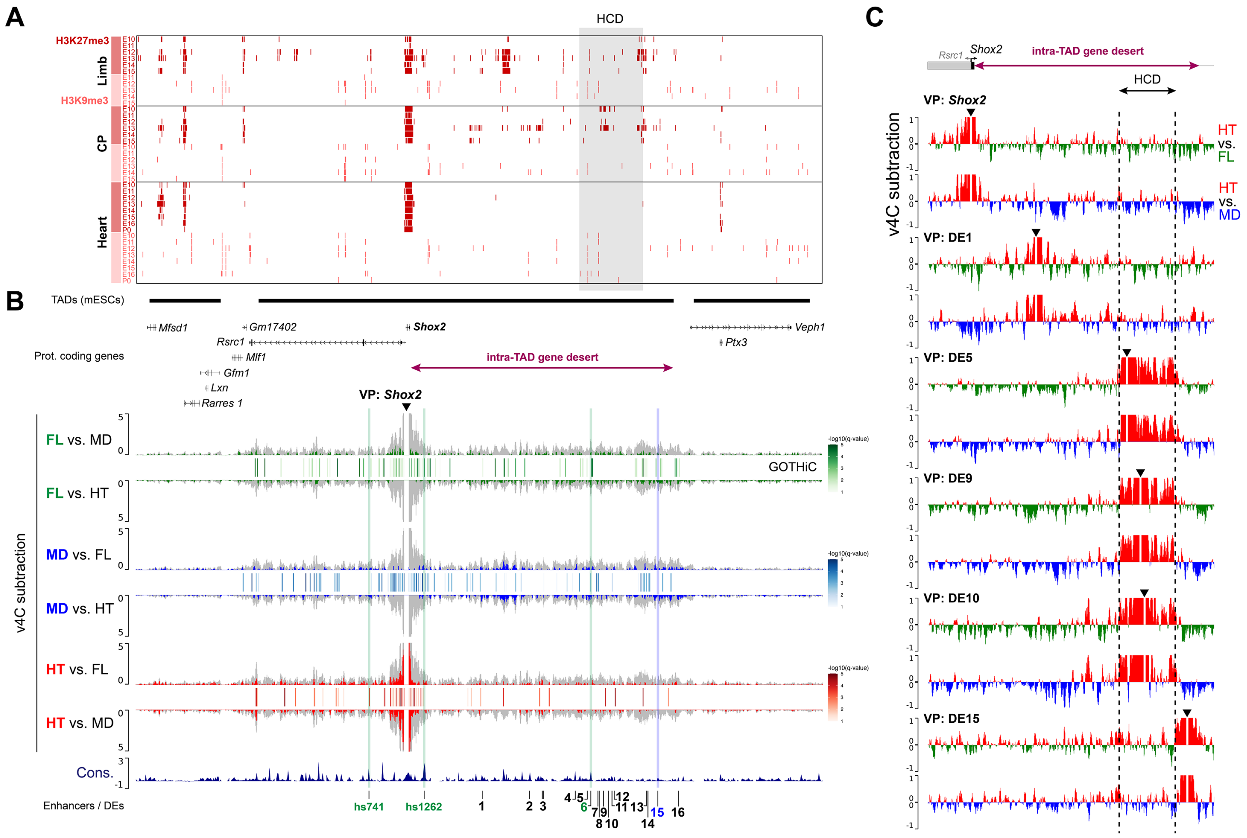


**Figure S3. Distribution of repressive histone marks and *Shox2* promoter contacts across the extended *Shox2* TAD.** (**A**) Distribution of ENCODE ChIP-seq peak calls from histone modification marks associated with repressive chromatin functions (H3K27me3, H3K9me3) in limb (E10.5-E15.5), craniofacial prominence (CP) (E10.5-E15.5) and heart tissues (E10.5-E16.5, P0)[2]. The high-density contact domain (HCD) interval (**Fig. 2**, **Fig. S2**) is shaded in gray. (**B**) Region-wide *Shox2* contacts based on virtual 4C (v4C) analysis from C-HiC in forelimb (FL), mandible (MD) and heart (HT) across the extended *Shox2* TAD (chr3:65977711-67631930) are shown in gray (*see also* **Fig. 2A**). V4C subtraction analysis indicates tissue-enriched contacts in pairwise comparisons (green: FL, blue: MD, red: HT). Bars between tissue-related profiles represent *Shox2*-contact regions corrected for bias and based on a significance threshold as identified by GOTHiC[4]. Validated enhancers with activities in the FL or MD are indicated by green and blue lines, respectively. TAD extensions from mESCs are shown[3]. Cons, Placental Conservation by PhyloP. 1-16, DEs. (**C**) V4C subtraction profiles of HT vs. FL and HT vs. MD comparisons using different VPs across the intra-TAD gene desert. VPs include *Shox2* and selected DEs (3kb intervals). In forelimb (green peaks) and mandible (blue peaks) the enhancer VPs within the HCD (DE5, 9 and 10) display numerous contacts with regions outside of the HCD, while in the heart (red peaks) tissue-enriched contacts from the HCD with outside regions appear reduced. VPs located outside of the HCD in the heart (DE 1, 15) show more distributed contacts.

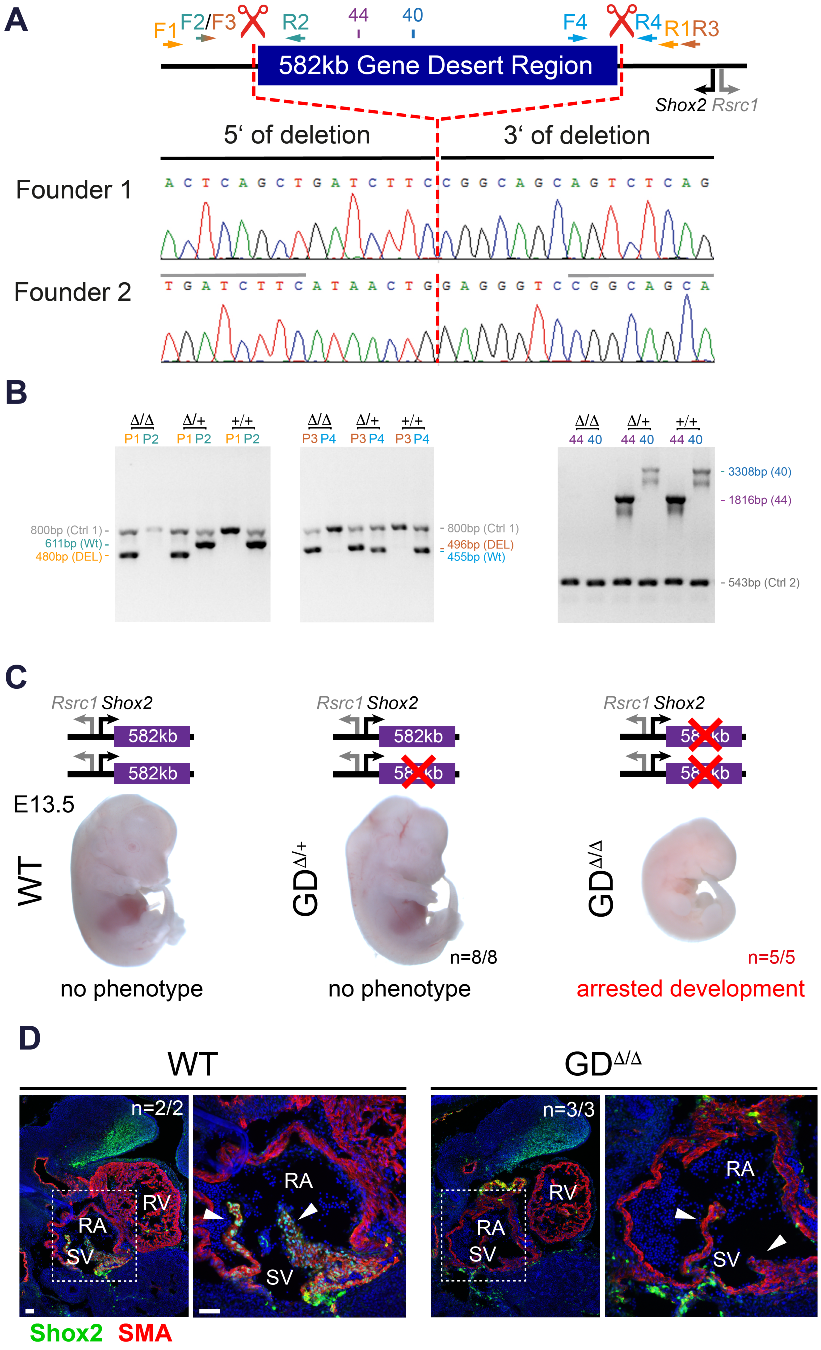


**Figure S4. Gene desert deletion triggers embryonic lethality linked to loss of *Shox2* in the SV.** (**A**) Sanger sequencing traces revealing clean gene desert deletion breakpoints (dashed red line) in the mouse lines generated via CRISPR-Cas9 for this study. Red scissors indicate the location of the guide RNAs used (**Table S4**). Location of primers (arrows) and amplicons (blocks) used for PCR are indicated (**Table S5**). (**B**) PCR validation and genotyping used to detect wildtype (+) and $GD^{\Delta}$ (DEL) alleles. Amplicon sizes are indicated on the side. Control primers (Ctrl-1 or Ctrl-2) amplifying an unrelated genomic region were used. P, primer pair (+ or DEL) mixed with control primers. (**C**) Homozygous gene desert deletion ($GD^{\Delta/\Delta}$) leads to arrested mouse embryonic development past E11.5, with full penetrance at E13.5, recapitulating the pattern observed in *Shox2*-deficient embryos[5]. Heterozygous ($GD^{\Delta/+}$) genotypes develop into viable and fertile mice. (**D**) Shox2 (green) is depleted in myocardial cells (SMA, red) of the SV including venous valves (white arrowheads) in

$GD^{\Delta/\Delta}$ embryos at E11.5. Nuclei are stained blue. Scale bars, 50μm. RA, right atrium. RV, right ventricle. SV, sinus venosus. N, number of biological replicates with similar results.

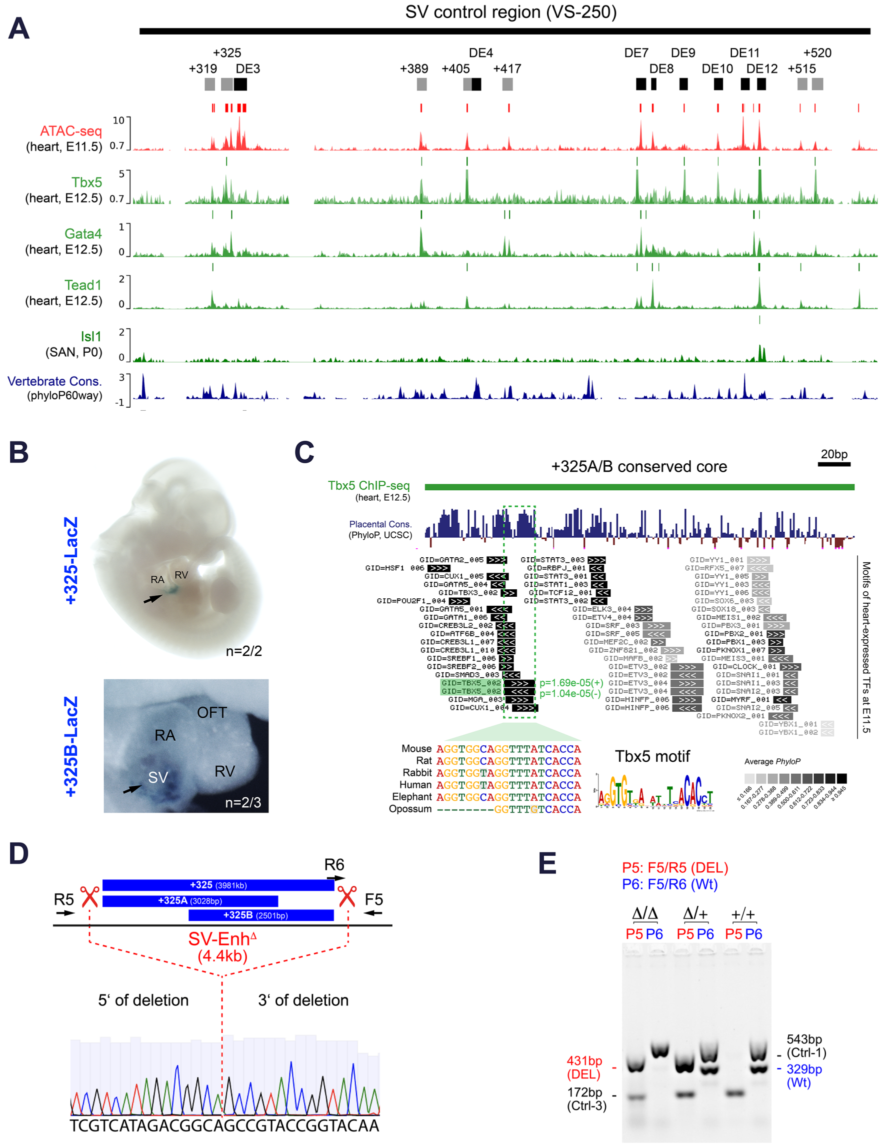


**Figure S5. TF binding profile and cardiac specificity of the +325 SV enhancer and its genomic deletion in mice.** (**A**) Reprocessed ChIP-seq profiles from cardiac TFs involved in SAN specification[6–8] (green) in the SV control region (VS-250)[9]. Open chromatin profiles (ATAC-seq) and vertebrate conservation track (Cons) are shown for comparison. Peak calls are indicated on top of each track. Genomic elements validated in this study are displayed on top (see also **Fig. 4D**). (**B**) Transgenic LacZ reporter validation of the +325 element (3981bp) and the +325B subregion (2501bp) at E11.5. N indicates transgenic replicates with similar staining. (**C**) Cardiac TF motif analysis (see ***Methods***) reveals a Tbx5 motif within the +325A/B conserved core region overlapping local Tbx5 enrichment in hearts at E12.5[6]. This motif is conserved across mammals and matches a TBX5 dimer DNA binding domain[10]. Cons., mammalian conservation by PhyloP (UCSC browser). All motifs annotated match a *p*-value of $\leq 10^{-4}$. *P*-values correspond to the probability of a random sequence (of the same length as the motif) producing a score that matches at least the one observed. Corresponding gene IDs (GID) are listed for each motif. (**D**) Sanger sequencing results of the deletion breakpoint in the +325 SV enhancer knockout allele (SV-Enh$^{\Delta}$) generated using CRISPR-EZ in mouse zygotes[11] (see ***Methods***). The location of primers used for PCR genotyping is indicated (**Table S5**). (**E**) PCR primer pairs (P5 and P6) to genotype the SV-Enh$^{\Delta}$ ($\Delta$) and wildtype (+) alleles were mixed with Ctrl-3 and Ctrl-1 primer pairs, respectively, to allow for amplification of a control band in the same PCR reaction.

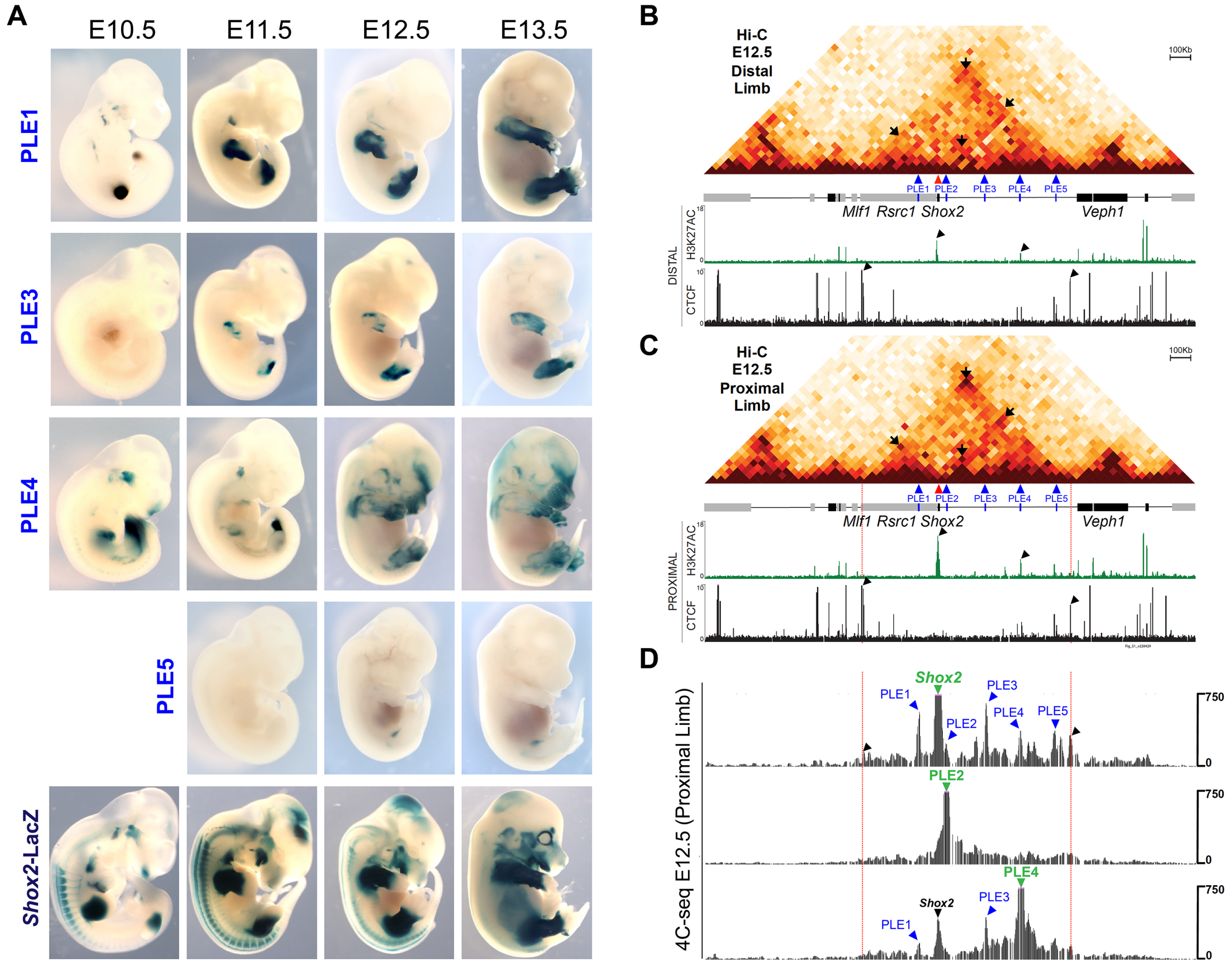


**Figure S6. Spatiotemporal activities of *Shox2*-PLEs and associated 3D-chromatin interactions.** (**A**) Developmental time course of proximal limb enhancer (PLE) activities in stable transgenic LacZ reporter lines compared to *Shox2* expression (*Shox2*-LacZ). For each element, one embryo from one representative transgenic line is shown per time-point (**Table S11** *and* **Methods**). PLE1 is the mouse ortholog of the hs741 enhancer[12,13]. To aid in visualization, embryos are depicted at progressively lower magnification at later stages. (**B, C**) Comparison of chromatin interactions within a 2.4Mb region centering the *Shox2* locus (mm10, chr3:65720000-68120000) in distal (**B**) and proximal (**C**) limbs at E12.5, including Hi-C, H3K27Ac and CTCF ChIP-seq tracks from a previous study[14]. *Shox2*-TAD boundaries are delineated by CTCF peaks (black arrowheads) and marked by red lines. Hi-C and H3K27ac profiles indicate that several interactions are stronger in proximal limb cells as compared to those in distal limb progenitors. Of the four highlighted interaction points (black arrows in the Hi-C map), two indicate *Shox2* interaction with U-dom and D-dom anchors (left and right arrows), while the upper arrow represents a strong interaction between TAD anchor points. The bottom arrow indicates a stronger interaction of *Shox2* with PLE3 in the proximal limb. Genes are shown as rectangles, with black indicating genes transcribed from left to right (telomeric to centromeric) and gray indicating genes transcribed in the opposite direction. (**C**) 4C-seq interaction profiles with PLE2 (hs1262/LHB-A) and PLE4 as viewpoints (purple arrowheads), compared to the profile based on the *Shox2* viewpoint (**Fig. 6B, Table S10**). One of two biologically independent replicates with similar results is shown.

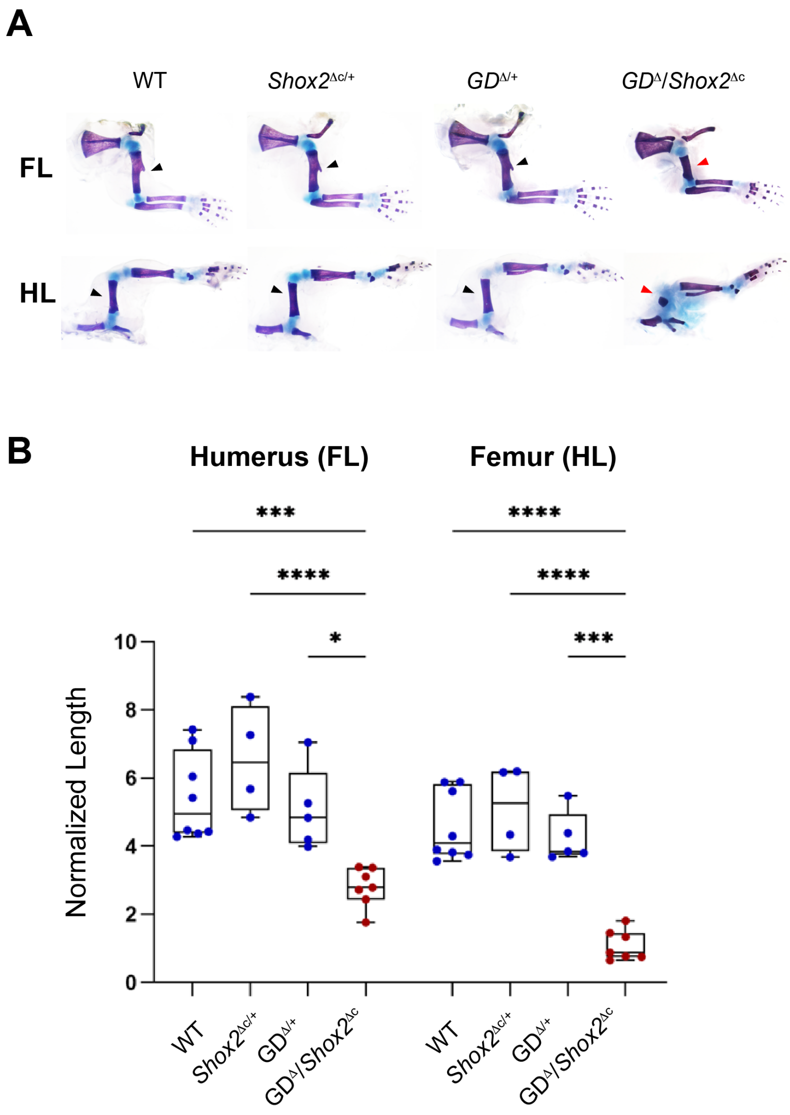


**Figure S7. Requirement of the gene desert for proximal limb development.** (**A**) The *Prx1*-Cre transgene was intercrossed with the conditional *Shox2*$^{\text{floxed/+}}$ allele to inactivate *Shox2* specifically in the mouse limb (*Shox2*$^{\Delta c}$). Preparation of limb skeletons from control (*Shox2*$^{\text{floxed/+}}$ [WT], *Shox2*$^{\Delta c/+}$, GD$^{\Delta/+}$) and sensitized gene desert knockout (GD$^{\Delta}$/*Shox2*$^{\Delta c}$) mice at P0 are shown. Stylopod elements in fore- and hindlimbs of GD$^{\Delta}$/*Shox2*$^{\Delta c}$ newborns are reduced in length (red arrowheads) compared to normal stylopod morphology in controls (black arrowheads). Chondrogenic skeletal elements are stained blue, ossified structures red. **(B)** Quantification of ossified humerus and femur compartments at P0 reveals a reduction in stylopod length in GD$^{\Delta}$/*Shox2*$^{\Delta c}$ mice compared to controls, similar to that found in adult limbs (**Fig. 6A, B**). Measurements were normalized to the length of the third metatarsal. Box plot indicates interquartile range, median, and maximum/minimum values (bars). Dots represent individual data points. ****, $P \leq 0.0001$; ***, $P \leq 0.001$; *, $P \leq 0.05$ (one-way ANOVA).

#### Supplementary Tables.

**Table S1 (provided as Excel file). Developmental enhancer predictions within the *Shox2* TAD.** *Sheet 1*: List of genomic elements within the extended *Shox2* TAD interval (chr3:65996078-67396078) that show significant ENCODE H3K27ac ChIP-seq enrichment in at least one tissue at any developmental stage and that were further filtered for ChromHMM[15] strong enhancers calls (as defined in Gorkin *et al.*, 2020). H3K27ac RPKM (Reads Per Kilobase of transcript, per Million mapped reads) values are shown for each developmental tissue and timepoint in a matrix format and are underlying the heatmap shown in **Figs. 1B and S1A**. For genes present in this domain, RNA-seq read counts are listed as fragments per kilobase of exon per million reads mapped (FPKM) (red shaded). Blue shades mark predicted gene desert enhancers (DEs). Mouse (mm10) coordinates (chrom, start, end) are given for each putative enhancer identified. Element IDs indicate distance to *Shox2* transcriptional start site (TSS). *Sheet 2*: The same matrix as in *sheet 1* but including q-values (-log10(q)) for each H3K27ac-enriched region as a measurement of statistical significance. *Sheet 3*: Metadata to be able to retrieve peak lists (for q-values) from the ENCODE Data Coordination Center (DCC, http://www.encodeproject.org/).

**Table S2. Primers used for PCR amplification of predicted desert enhancer (DE) elements for Hsp68-*LacZ* reporter assays.** Distance (in *kb*) from the *Shox2* TSS is indicated in brackets for each element (-, upstream; +, downstream).

| *Element ID* | *Forward primer* | *Reverse primer* | *Product Size (bp)* | *Genomic coordinates (mm10)* |
|---|---|---|---|---|
| **DE1 (+183)**<br>(mm1849) | TCCAACTAGCCACAATCCACTA | GGTTGACAAAGGTTCAGAAAGG | 2486 | chr3<br>66797249<br>66799734 |
| **DE2 (+297)**<br>(mm1852) | TCCTCTCTGTGTTTCAGCTTTG | TGGGTGACTCAGGTAAACCTCT | 3167 | chr3<br>66682968<br>66686134 |
| **DE3 (+330)**<br>(mm1853) | ACCATGGTAGGAAGTTCATTGG | GTTAGAGCTGTTGGGAAAATGC | 4408 | chr3<br>66650021<br>66654428 |
| **DE4 (+408)**<br>(mm1837) | GCTATACGCCGTCAGCTTTAGT | ATGTGAATGAAGCACAAATTGC | 3236 | chr3<br>66572737<br>66575972 |
| **DE5 (+437)**<br>(mm2108) | GATGTGGGGAAACTTCTGAAAC | TACAGACCCAGACAAGAGCAGA | 4335 | chr3<br>66542058<br>66546392 |
| **DE6 (+444)**<br>(mm1845) | GATGCAGGCACGATATACAAAA | AGACCTTACACACGTGCACAAC | 2962 | chr3<br>66535471<br>66538432 |
| **DE7 (+463)**<br>(mm2103) | CTGCGCTTTCTTCTTATCCCTA | CAGATCCACCTCTTCCTTCATC | 3402 | chr3<br>66518263<br>66521664 |
| **DE8 (+466)**<br>(mm1838) | GGAATTGCTTTGTAGCTCTGCT | CAGGGAGGAAGCTTCTAGTTCA | 1816 | chr3<br>66514951<br>66516766 |
| **DE9 (+475)**<br>(mm1846) | GACACCACCAAGAGTTCGTGTA | AATTACAATGTGTGGGGGAGAC | 2824 | chr3<br>66504602<br>66507425 |
| **DE10 (+487)**<br>(mm2109) | TCTCTATGACCAAACGGGCTAT | GGATTTGGAAGAACAAGAGGTG | 3109 | chr3<br>66492941<br>66496049 |
| **DE11 (+495)**<br>(mm2110) | CTGTGTATGCCTTTGCTCTCAG | CTCTGCTCATATTCTGCCTCCT | 3067 | chr3<br>66484145<br>66487211 |
| **DE12 (+501)**<br>(mm1839) | CTGCTCTAATTCTGGGAGGTTG | TTATTGCTTGGTGAGAATGTGG | 3041 | chr3<br>66478804<br>66481844 |
| **DE13 (+581)**<br>(mm1842) | TGTATTCCACAGCCTCCCTAGT | CCCAAGGTCTGGTTTAGAACTG | 2511 | chr3<br>66400179<br>66402689 |
| **DE14 (+583)**<br>(mm2111) | TCCTACAGGCAAGACCTCTCTC | CATGGTCCAACTGGTATTGATG | 1942 | chr3<br>66397740<br>66399681 |
| **DE15 (+606)**<br>(mm1843) | CATTGGTACTTGGGCTGAAAA | TTACAAAGCTCCTGACGCAGT | 3139 | chr3<br>66373217<br>66376355 |
| **DE16 (+656)**<br>(mm2112) | CAGAGGTCCTGAACTCAATTCC | TCCTGCTGTGCATAGAACAACT | 2839 | chr3<br>66324764<br>66327602 |

**Table S3 (provided as Excel file). C-HiC domain boundaries and *Shox2*-interacting regions by GOTHiC.** *Sheet 1*: Genomic coordinates (mm10) of inter-domain boundaries, related p-values and TAD separation scores (TAD_SEP) resulting from HiCExplorer[16] analysis of *Shox2* C-HiC datasets (interrogated interval: chr3:65196078-68696078). $P<10^{-12}$ is listed as “0”. FL, forelimb. MD, mandible. HT, Heart. *Sheets 2-4*: Genomic coordinates (mm10) of elements showing significant Hi-C contacts with *Shox2* as determined by GOTHiC[4] using a 10kb viewpoint on the *Shox2* TSS in forelimb (FL), mandible (MD) or heart (HT) tissues (interrogated interval: chr3:65196078-68696078), respectively.

**Table S4. CRISPR deletions and sgRNA templates**. Genomic coordinates of the CRISPR-deleted regions are provided for each founder mouse line. For the gene desert deletion, use of unique sgRNAs resulted in the generation of two nearly identical founder lines (see **Fig. S3A, B**).

| Mouse allele | Genomic coordinates of deletion (mm10) | Deleted region (bp) | 5' sgRNA target sequence | 5' sgRNA target sequence |
|---|---|---|---|---|
| **GD$^{\Delta}$** (Founder 1) | chr3 66365062 66947168 | 582107 | TGATCTTCATAACTGCCATGGGG | TGAAGCACAAGGCTGGCGGGAGG |
| **GD$^{\Delta}$** (Founder 2) | chr3 66365069 66947161 | 582093 | TGATCTTCATAACTGCCATGGGG | TGAAGCACAAGGCTGGCGGGAGG |
| **SV-Enh$^{\Delta}$** | chr3 66654441 66658882 | 4442 | GGGATACATTGAGACCGGCA | ACAGCAGTATCTGCCGTAGA |

**Table S5. Primers used for screening and genotyping of CRISPR deletion mouse strains.** PCR genotyping strategy and results using agarose gel electrophoresis are shown in **Fig. S4A, B** (for the gene desert deletion) and **S5D, E** (for +325 SV enhancer deletion). GD, gene desert. Del, deletion. Enh, enhancer. P, primer pair. f, founder. N.A., not amplified.

| *Analyzed Region* | *Primer name* | *Sequence* | *Product Size (bp)* |
|---|---|---|---|
| **GD-del (P1)** | **F1** | agcggagggatactttagcac | WT: 582587 (N.A.) |
| | **R1** | tgctgagagatgaaccctgat | KO: 480 (f1) / 494 (f2) |
| **GD-del 5' junction (P2)** | **F2** | ccgcagagttctttgagagttt | WT: 611 |
| | **R2** | gacccagcagtattcggagtta | KO: N.A. |
| **GD-del (P3)** | **F3** | ccgcagagttctttgagagttt | WT: 582603 (N.A.) |
| | **R3** | acaagagcatgtgttcaagtgg | KO: 496 (f1) / 510 (f2) |
| **GD-del 3' junction (P4)** | **F4** | tgccctacagaagttaagcaca | WT: 455 |
| | **R4** | tactgttgccatcactccattc | KO: N.A. |
| **Region 44 (+466kb)** | **44 F** | ggaattgctttgtagctctgct | WT: 1816 |
| | **44 R** | cagggaggaagcttctagttca | KO: N.A. |
| **Region 40 (+389kb)** | **40 F** | tctataacggagctgcacttga | WT: 3308 |
| | **40 R** | ggcatttgtgagacatgagaaa | KO: N.A. |
| **SV-Enh-del (P5)** | **F5** | ccaggataggaagaaagcaaga | WT: 4873 (N.A.) |
| | **R5** | agcaaggaggacaccaaagtag | KO: 431 |
| **SV-Enh 5' junction (P6)** | **F5** | ccaggataggaagaaagcaaga | WT: 329 |
| | **R6** | taaagctagtccgtctcgtgtg | KO: N.A. |
| **Control region 1 (Ctrl-1)** | **Ctrl-1 F** | ccctagttctgtaaaccaggcta | WT/KO: 800 |
| | **Ctrl-1 R** | tcatgtgtcttaggagagggttc | (Tbx3 locus) |
| **Control region 2 (Ctrl-2)** | **Ctrl-2 F** | agctggtagccttaaaataagccaa | WT/KO: 543 |
| | **Ctrl-2 R** | gcctgaaagaggtcatcatcacc | (Gli3 locus) |
| **Control region 3 (Ctrl-3)** | **Ctrl-3 F** | ggaatgcccaggacataaaa | WT/KO: 172 |
| | **Ctrl-3 R** | gagaagggcttatttcagctcac | (Gata4 locus) |

**Table S6. Primers used for SYBR Green Real-time PCR analysis.**

| Target Gene | qPCR primer | Sequence | *Product Size (bp)* |
|---|---|---|---|
| ***Shox2*** | Shox2_F | CCCGAGTACAGGTTTGGTTTC | 119* |
| | Shox2_R | GAAGCTTGTAGAGTTGCACCC | |
| ***Rsrc1*** | Rsrc1_F | TGCAATTGGTCCTTGAAGCT | 104* |
| | Rsrc1_R | GGTGGCTTGGTCTTCTTCTT | |
| ***Actb*** | Actb_F | ACACTGTGCCCATCTACGAGG | 280* |
| | Actb_R | CATCACTATTGGCAACGAGCG | |

*primer pair validated and used in a previous study[24].

**Table S7 (provided as Excel file). Reprocessed ATAC-seq and ChIP-seq datasets used in this study.** List of previously published mouse ATAC-seq and ChIP-seq datasets from embryonic hearts (E12.5), sinoatrial node (SAN, P0) and limb re-processed as part of this study using uniform pipelines (see **Methods**). The resulting NarrowPeak (NP) and peak intersection (Int) datasets are indicated and provided in **Supplementary Data File 1**. FL, forelimb. HL, hindlimb. r, replicate.

**Table S8. Genomic elements tested via transgenic LacZ reporter assays based on cardiac ATAC-seq enhancer predictions.** All elements were validated using conventional random transgenesis based on Hsp68-LacZ reporter constructs[17], except for the +325 element which was analyzed using site-directed transgenesis at the *H11* safe-harbor locus based on a targeting vector encoding a LacZ reporter unit with a minimal β-globin promoter[18,19]. Distance (in *kb*) from the *Shox2* TSS and Vista IDs are indicated in brackets for each element (-, upstream; +, downstream).

| *Element ID* | *Forward primer* | *Reverse primer* | *Product Size (bp)* | *Genomic coordinates (mm10)* |
|---|---|---|---|---|
| **+319 (mm2105)** | GGTCAGGAATTCAGAGGTCAAC | ATACATCTGGGTTTGTCCATCC | 3463 | chr3 66660575 66664037 |
| **+325** | ATCAGCTCAGCTTTGGTTAAGG | ACTGACCCTTCACAGACTGGTT | 3982 | chr3 66654695 66658676 |
| **+325-A (mm2106)** | GCCATTATGGTCTTGAAGGAAG | ACTGACCCTTCACAGACTGGTT | 3029 | chr3 66655648 66658676 |
| **+325-B (mm2114)** | ATCAGCTCAGCTTTGGTTAAGG | GAATTCCTGATGCACTCTTTCC | 2502 | chr3 66654695 66657196 |
| **+389 (mm2099)** | TCTATAACGGAGCTGCACTTGA | GGCATTTGTGAGACATGAGAAA | 3307 | chr3 66590716 66594023 |
| **+405 (mm2102)** | GATGTGGGGAAACTTCTGAAAC | TACAGACCCAGACAAGAGCAGA | 3036 | chr3 66575695 66578731 |
| **+417 (mm2101)** | CTGCCATAACATTTGTGCTGTT | AATGCTTGTTTCCCAGAAGGTA | 3540 | chr3 66562316 66565856 |
| **+515 (m2100)** | GGTTGACACAAGTAACCAGCAA | GCAAGCACTCTACCCCATAATC | 3335 | chr3 66465121 66468456 |
| **+520 (mm2115)** | GTATGTTGTGGGCTTTCTCCTC | ATGAATCCCATGTAAGCAAACC | 3846 | chr3 66459872 66463718 |

**Table S9 (provided as Excel file). List of PWMs from heart-expressed TFs for genome-wide cardiac TF motif prediction.** Of the 1’376 motifs obtained, 14 (shaded in gray) were part of larger motifs and thus omitted in the analysis (see **Methods**).

**Table S10. Viewpoints and primers used for 4C-Seq.**

| **Viewpoint** | **Genomic coordinates** (NlaIII fragment) (mm10) | **Primer sequence:**<br>Illumina adapter sequences are shown in italics.<br>Sequence specific to the viewpoint in bold. |
|---|---|---|
| ***Shox2*** | chr3:66,980,317-66,981,259* | Forward/Reading primer:<br>*AATGATACGGCGACCACCGAACACTCTTTCCCTACACGACGCTCTTCCGATCT*<br>**CCAATTAAGAAAATATGTGGCATG**<br><br>Reverse Primer: *CAAGCAGAAGACGGCATACGA***AGAATGTGAAGTTTGGTCCC** |
| **PLE2** | chr3:66,938,480-66,939,521 | Forward/Reading primer:<br>*AATGATACGGCGACCACCGAACACTCTTTCCCTACACGACGCTCTTCCGATCT*<br>**ACTGCTTAGTAAAGACTAATTATTCATG**<br><br>Reverse Primer:<br>*CAAGCAGAAGACGGCATACGA***AATGACATTATTATAAAATGCAATACTCT** |
| **PLE4** | chr3:66,573,586-66,574,775 | Forward/Reading primer:<br>*AATGATACGGCGACCACCGAACACTCTTTCCCTACACGACGCTCTTCCGATCT*<br>**GGCTGATTCTCCTGCATG**<br><br>Reverse Primer:<br>*CAAGCAGAAGACGGCATACGA***AGTTATAAAGATGATTAAGCTCTGATC** |

*The *Shox2* viewpoint spans the 3' end of the first *Shox2* exon and the 5' end of intron 1.

**Table S11. Proximal limb enhancers (PLEs) identified via 4C-seq and validated by *β*-globin promoter-*LacZ* transgenesis.** Distance (in *kb*) from the *Shox2* TSS is indicated in brackets for each element (-, upstream; +, downstream).

| *Element ID* | *Forward primer* | *Reverse primer* | *Product Size (bp)* | *Genomic coordinates (mm10)* |
|---|---|---|---|---|
| **PLE1 (-89)** | TGGGCAAAGATCACAGAACA | GTGTGTGTGTGTGTGGTGGA | 1674 | chr3:67070163-67071836 |
| **PLE2 (+43)** | GAAGGACCGCACAGCTTATC | GGTCCACATATGCCCAAGGA | 2428 | chr3:66937659-66940086 |
| **PLE3 (+237)** | GAAGAGGGGGCAGATTGTGTTGACTG | TGCTTCTTCAAATATTGCTTTGCTAAT | 10351* | chr3:66739935-66750285 |
| **PLE4 (+407)**** | GTGAATGAAGCACAAATTGCAA | AAAGCCCATGTGTTCATCCCAG | 3718 | chr3:66572253-66575970 |
| **PLE5 (+568)** | GGTCTATCTTGTTGCATGTTTTGTT | GGACAAACAGAGCTCAGAAGAGA | 9473*** | chr3:66409729-66419201 |

*A 9128bp ApaI/SalI sub-fragment (mm10: chr3:66740432-66749559) of the 10351bp PCR fragment was cloned into the pβlacz vector and used for *LacZ* transgenesis.
**The PLE4 fragment contains the DE4 element (**Table S2**) and an additional 486bp.
***A 8520bp ApaI/SalI sub-fragment (mm10: chr3:66409729-66418248) of the 9473bp PCR fragment was cloned into the pβlacz vector and used for *LacZ* transgenesis.

### SUPPLEMENTARY REFERENCES


1. Dixon, J. R. *et al.* Topological domains in mammalian genomes identified by analysis of chromatin interactions. *Nature* **485**, 376–380 (2012).
2. Gorkin, D. U. *et al.* An atlas of dynamic chromatin landscapes in mouse fetal development. *Nature* **583**, 744–751 (2020).
3. Bonev, B. *et al.* Multiscale 3D Genome Rewiring during Mouse Neural Development. *Cell* **171**, 557-572.e24 (2017).
4. Mifsud, B. *et al.* GOTHiC, a probabilistic model to resolve complex biases and to identify real interactions in Hi-C data. *PLoS One* **12**, e0174744 (2017).
5. Blaschke, R. J. *et al.* Targeted mutation reveals essential functions of the homeodomain transcription factor Shox2 in sinoatrial and pacemaking development. *Circulation* **115**, 1830–1838 (2007).
6. Akerberg, B. N. *et al.* A reference map of murine cardiac transcription factor chromatin occupancy identifies dynamic and conserved enhancers. *Nat. Commun.* **10**, 4907 (2019).
7. He, A. *et al.* Dynamic GATA4 enhancers shape the chromatin landscape central to heart development and disease. *Nat. Commun.* **5**, 4907 (2014).
8. Liang, X. *et al.* Transcription factor ISL1 is essential for pacemaker development and function. *J. Clin. Invest.* **125**, 3256–3268 (2015).
9. van Eif, V. W. W. *et al.* Genome-Wide Analysis Identifies an Essential Human TBX3 Pacemaker Enhancer. *Circ. Res.* **127**, 1522–1535 (2020).
10. Jolma, A. *et al.* DNA-binding specificities of human transcription factors. *Cell* **152**, 327–339 (2013).
11. Chen, S., Lee, B., Lee, A. Y.-F., Modzelewski, A. J. & He, L. Highly Efficient Mouse Genome Editing by CRISPR Ribonucleoprotein Electroporation of Zygotes. *J. Biol. Chem.* **291**, 14457–14467 (2016).
12. Ye, W. *et al.* A unique stylopod patterning mechanism by Shox2-controlled osteogenesis. *Development* **143**, 2548–2560 (2016).

13. Osterwalder, M. *et al.* Enhancer redundancy provides phenotypic robustness in mammalian development. *Nature* **554**, 239–243 (2018).

14. Rodríguez-Carballo, E., Lopez-Delisle, L., Yakushiji-Kaminatsui, N., Ullate-Agote, A. & Duboule, D. Impact of genome architecture on the functional activation and repression of Hox regulatory landscapes. *BMC Biol.* **17**, 55 (2019).

15. Ernst, J. & Kellis, M. ChromHMM: automating chromatin-state discovery and characterization. *Nat. Methods* **9**, 215–216 (2012).

16. Wolff, J., Backofen, R. & Grüning, B. Loop detection using Hi-C data with HiCExplorer. *Gigascience* **11**, (2022).

17. Osterwalder, M. *et al.* Characterization of Mammalian In Vivo Enhancers Using Mouse Transgenesis and CRISPR Genome Editing. *Methods Mol. Biol.* **2403**, 147–186 (2022).

18. Kvon, E. Z. *et al.* Comprehensive In Vivo Interrogation Reveals Phenotypic Impact of Human Enhancer Variants. *Cell* **180**, 1262-1271.e15 (2020).

19. Darbellay, F. *et al.* Chondrogenic Enhancer Landscape of Limb and Axial Skeleton Development. *bioRxiv* 2023.05.10.539849 (2023) doi:10.1101/2023.05.10.539849.
